# Selection and characterization of DNA aptamers targeting the surface borrelial protein CspZ with high-throughput cross-over SELEX

**DOI:** 10.1101/2025.01.13.632687

**Authors:** Mickaël Guérin, Marylène Vandevenne, André Matagne, Willy Aucher, Julien Verdon, Emmeline Paoli, Jules Ducrotoy, Stéphane Octave, Bérangère Avalle, Irene Maffucci, Séverine Padiolleau-Lefèvre

## Abstract

Lyme borreliosis (LB) is the most prevalent tick-borne illness, with an estimated 700 000 cases annually in the United States and Europe. LB diagnosis based on two-tiered serology remains controversial due to its indirect nature and low sensitivity during the early stage of the disease. Aptamers are single-stranded DNA or RNA oligonucleotides that exhibit high selectivity and specificity for their target due to their unique three-dimensional structure. By applying cross-over-SELEX process, an enrichment of DNA oligonucleotide sequences against a surface protein of *Borrelia*, named CspZ, has been performed and monitored using absorbance at 260 nm, melting curves and NGS analyses. Beyond sequence enrichment, oligonucleotides binding to CspZ were observed during the selection rounds by Dot Blot and beads assays. Thirteen unique and highly redundant oligonucleotide sequences have been further characterized using multiple approaches such as Dot Blot, BioLayer Interferometry and Surface Plasmon Resonance. The selected aptamers showed K_D_ values from tens of nanomolar to the micromolar range by BLI and SPR. Two aptamers, characterized by flow cytometry and epifluorescence microscopy, were able to specifically recognize *Borrelia burgdorferi* sensu stricto. This novel strategy holds promise for the development of an improved diagnosis assay.

**GRAPHICAL ABSTRACT:** 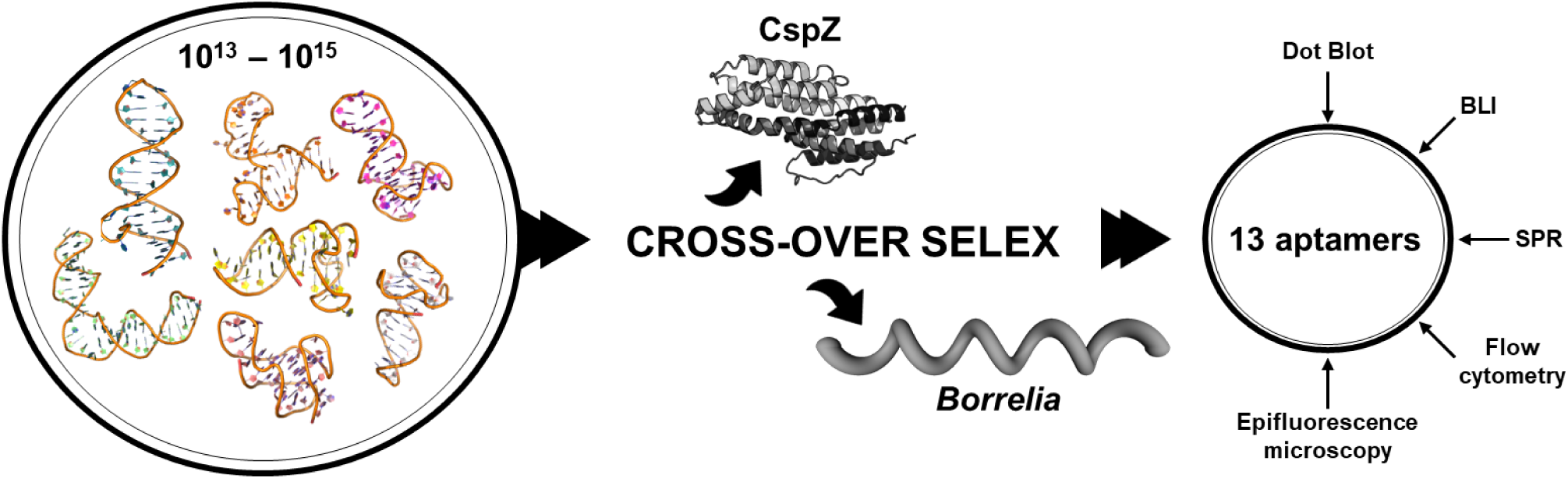

**HIGHLIGHTS:**

- DNA aptamers were selected by cross-over SELEX.
- Aptamers’ KDs ranged from tens of nanomolar to the micromolar range.
- Aptamer characterization was performed by Dot Blot, beads assay, BLI, and SPR.
- Interaction with *Borrelia* was tested by flow cytometry and epifluorescence microscopy.
- Aptamers bind to both the recombinant and borrelial surface CspZ protein.

## INTRODUCTION

Lyme borreliosis (LB), transmitted by ticks of the genus *Ixodes* and caused by the complex *Borrelia burgdorferi* (*Bb*) sensu lato (sl), is the most common vector-borne infectious disease in the United States and Europe, with more than 700000 estimated cases each year. To date, LB diagnosis relies either on the presence and identification of a pathognomonic skin manifestation, called Erythema migrans, or on an indirect diagnosis. This latter is based on a two-tiered standard serology, an indirect approach which suffers from a low sensitivity depending on the stage of the disease (1–3). Many efforts have been joint to enhance sensitivity and rapidity of tests, leading to the reporting of new probes or new methods, such as ELISpot, phage-PCR or nucleic acid aptamer sensors (4–9).

Nucleic acid aptamers are single-stranded DNA or RNA oligonucleotides capable of specifically binding to a wide range of molecules with dissociation constants in the range of nano- to picomolar. Aptamers have been shown to have a plethora of applications, such as biosensing, diagnosis and therapy (10–12), including the detection of biotoxins, pesticides, and pathogenic microorganisms (13). Notably, they have been successfully utilized to detect various biomarkers in complex biological samples such as blood (14) and urine (15). The specific binding properties of aptamers arise from their original and unique three-dimensional structures (16–18), which allow specific non-covalent interactions with the target (13, 19). Aptamers also display minimal batch-to-batch variability and can be produced at large scale. Moreover, they exhibit enhanced chemical and thermal stability as compared to other types of biomolecules, therefore facilitating extended storage and handling procedures. Their low molecular weight, which allows for rapid clearance from the bloodstream without organ accumulation, coupled with their non-toxic nature, minimal immunogenicity, and cost-effective synthesis, further contribute to their attractiveness (11, 20–23). Aptamers are commonly selected through an *in vitro* process known as **S**ystematic **E**volution of **L**igands by **EX**ponential enrichment (SELEX), first described in 1990 by Tuerk and Gold (24) and Ellington and Szostak (25). SELEX technology has undergone significant advancements over the years, leading to the development of different variants, such as capillary electrophoresis (CE)-SELEX, magnetic bead-based SELEX (sometimes combined with fluorescence, termed FluMag-SELEX), and Cell-SELEX (26).

With the aim of conducting SELEX-based evolution of aptamers recognizing the spirochetes responsible for Lyme borreliosis (LB), we focused our interest on one borrelial protein called CspZ, also known as BbCRASP-2/BBH06, a membrane lipoprotein exposed to the outer bacterial surface. While its surface expression decreases during *in vitro* cultivation, CspZ remains abundantly produced during infection in mammalian cells (27). Thus, this protein is a promising candidate for the conceptualisation of a direct detection towards it and, therefore, for LB diagnosis. Additionally, the *cspZ* gene, positioned on the linear plasmid lp28-3 of *Bb* B31 strain, is highly conserved with about 80% identity among the subspecies responsible for LB (*B. afzelii, B. garinii, B. bavariensis,* and *B. spielmanii*), making the product protein CspZ a common biomarker for several *Borrelia* strains. Furthermore, previous biochemical and biophysical characterization of CspZ interactions with human complement factor H and complement factor H like-1 (28–31), coupled with resolved structural data of the complexes (PDB: 6ATG) (32) prove the existence of accessible epitopes on the cell surface.

Cross-over SELEX offers a synergistic combination of the advantages inherent to FluMag SELEX and cell-SELEX. While it inherits the simplicity, low cost, and ease of application from FluMag SELEX, it also shares the ability of cell-SELEX to select aptamers against accessible epitopes in their native conformation (33). Furthermore, it mitigates many of the drawbacks associated with these individual methods. Finally, while counter-selection using knockout bacteria is crucial in cell-SELEX to eliminate non-specific binding, cross-over SELEX can succeed without it, thanks to the combination with FluMag-SELEX, which involves alternating protein capture and negative selection.

In this study, we aimed to develop new DNA aptamers against CspZ, by using a cross-over SELEX approach (combination of FluMag-SELEX and, in this study, a single round of cell-SELEX). This approach allowed for the initial enrichment of CspZ-binding aptamers, followed by a selection round using whole bacterial cells to influence the selection of aptamers targeting exposed epitopes in their native environment. We propose a proof of concept for the development of new direct probes of borrelial pathogens involved in infectious diseases. To do so, we monitored the SELEX process and the progressive convergence of oligonucleotide diversity towards CspZ. After detailed inspection through Next-Generation Sequencing (NGS), we evaluated the affinity and selectivity of each selected aptamer with integrative molecular approaches regrouping Dot Blot analysis, biolayer interferometry (BLI) and surface plasmon resonance (SPR). Then, after determining the optimal CspZ expression time point through Reverse Transcription-quantitative polymerase chain reaction (RT-qPCR) analysis, the capacity of selected aptamers to bind to the bacteria was evaluated using epifluorescence microscopy and flow cytometry.

## MATERIAL AND METHODS

### Materials and instruments

All chemicals for the preparation of buffers, solutions and materials were purchased from VWR International (Germany), Merck (Germany) or Thermo Fisher (USA). The HisPur™ Ni-NTA Magnetic Beads kit was purchased from Thermo Scientific (USA), whereas the Glutathione High Capacity Magnetic Agarose Beads kit was purchased from Merck. Regarding PCR components, dNTP and Monarch® PCR & DNA Cleanup Kit (5 μg) were purchased from New England Biolabs (NEB, USA), and NeoProof DNA polymerase and Neogreen qPCR Master Mix (2x), were from CliniSciences (France).

The concentration measurements of the DNA were performed using NanoDrop™ 2000 and the fluorescence measurements were performed on the VANTAstar® microplate reader from BMG labtech (Germany). Sodium Dodecyl-Sulfate - PolyAcrylamide Gel Electrophoresis (SDS-PAGE), Western-Blot (WB) and Dot-Blot pictures were taken with the ChemiDoc™ imaging system (Biorad, USA). PCR and melting curve analysis have been performed in a C1000 Touch thermal cycler and in a Chromo4 Four-Color Real-Time PCR Detection System (Biorad, USA), respectively. All BLI experiments were performed on the OCTET HTX instrument (ForteBio, Sartorius, Gottingen, Germany) at the Robotein facility, which is part of the BE-Instruct ERIC Center. Protein-DNA interactions were also analysed by SPR technique using a Biacore T100 instrument (Cytiva). To assess the expression of *cspZ*, qPCR analysis was performed on a LightCycler 480 (Roche). To evaluate the ability of selected aptamers to bind available epitopes on bacteria, flow cytometry experiments were performed using a CytoFlex flow cytometer (Beckman Coulter Life Sciences, Indianapolis, USA) and the CytExpert software. *Bb* B31 was purchased from ATCC.

The SELEX library was purchased from Eurogentec (Belgium). The random 76-mer ssDNA library comprised a central random region of 40 nucleotides (N40) flanked by 18 nucleotides primer binding sites for amplification and cloning: 5’-ATACCAGCTTATTCAATT-N40-AGATAGTAAGTGCAATCT-3’. The composition of the library has been designed by requiring 25% adenine (A), 25% thymine (T), 25% cytosine (C) and 25% guanine (G). The library was purified by PAGE. The following primers, also purchased from Eurogentec, were used either for the amplification during SELEX or for the final amplification step before cloning: 5’-ATACCAGCTTATTCAATT-3’ with and without 5’-6-Carboxyfluorescein (FAM) and 5’-AGATTGCACTTACTATCT-3’ with and without 5’-poly-A20-hexaethylene glycol spacer (HEGL). All the primers were purified by Reverse Phase - High-Pressure Liquid Chromatography (RP-HPLC). The synthetic ssDNA library and the primers were dissolved in nuclease-free water to a final concentration of 100 µM. For screening analysis, all the oligonucleotides were purchased from Eurofins Genomics (Luxembourg) and purified using HPLC. For characterization studies of aptamers, the oligonucleotides were purchased from Eurogentec using PAGE purification. Their sequences are listed in the **Supplementary Table 1**, including a random 76-nt sequence (AptaCtrl) with equal content of each nucleotide and reported in NGS sequencing of the initial library but not in following rounds of selection, used as a negative control.

### Production and purification of target protein (CspZ) and its protein functional mimic (FhbA)

Three plasmids containing the *cspZ* gene from *Bb*ss (NCBI:txid139) were used. Two plasmids coding for CspZ with a N-terminal 1.7-kDa hexahistidine tag (6His-CspZ) were kindly provided by Prof. Marconi (pET-45b(+)) and by Dr Brangulis (pETM-11). The third plasmid coding for Glutathione S-Transferase (GST) – CspZ fusion protein was kindly provided by Prof. Kraiczy (pGEX-6P-1). Additionally, the plasmid for 6His-FhbA from *B. hermsii* (NCBI:txid140) was kindly provided by Dr Kogan (pET-32 Ek/LIC). *E. coli* BL21 (DE3)-chemical competent cells were transformed with each plasmid by heat-shock at 42 °C for 45 seconds. Protein expression was induced by addition of 1 mM isopropyl-β-d-thiogalactopyranoside (IPTG) for 3 h, and the cells were harvested by centrifugation (6, 000 × g, 10 min) and lysed using B-PER™ Complete Bacterial Protein Extraction Reagent (reference 78243, Thermofisher) supplemented with a cOmplete™ ULTRA EDTA-free protease inhibitor tablet (reference 5892953001, Roche Diagnostics).

Purification of recombinant proteins 6His-CspZ and 6His-FhbA were performed as described in Guérin and co-workers (31). Regarding the GST fused-CspZ, the recombinant proteins were purified from clarified cell lysate by glutathione (GSH) affinity chromatography using a 1 mL HiTrap® TALON® Crude column. After washing with PBS, the retained proteins were eluted by increasing concentrations of reduced GSH from 0 to 40 mM in PBS. After dialysis in PBS, recombinant protein concentrations were measured by Bicinchoninic Acid (BCA) or Pierce 660 nm kits respectively. The purity was verified by SDS-PAGE, Western blot and HPLC (31). Briefly, for HPLC analysis, an UltiMate 3000 HPLC from Thermofisher with a Fortis Bio C18, 3 µm, 4.6 × 100 mm reversed phase column was used. Two mobile phases (1) ultrapure water with 0.1 vol % Trifluoroacetic acid (TFA) and (2) acetonitrile (ACN) with 0.1 vol % TFA were used for the linear gradient elution between 5% to 100% ACN in 10 minutes. The solvent flow rate was set at 1 mL/min. Quantification was performed using external calibration peak area measurement.

### Coating of the target protein on magnetic beads for the aptamer’s selection

Saturation of beads surface was performed using 200 µg of CspZ, either conjugated to His Tag or fused to GST, immobilized on each type of beads (160 µL of beads solution were used for HisPur™ Ni-NTA Magnetic Beads, whereas 50 µL were used for the Glutathione High Capacity Magnetic Agarose Beads). Briefly, 400 µg of CspZ were diluted from stock in 1.2 mL of PBS buffer and incubated for 1 hour on a rotating wheel. The protein-coated beads were separated using a magnetic rack and washed twice with 1 mL of Binding/Wash (B/W) buffer (100 mM NaCl, 20 mM Tris-HCl pH 7.6, 2 mM MgCl_2_, 5 mM KCl, 1 mM CaCl_2_, 0.02% Tween 20).

### Flu-Mag SELEX rounds of selection

FluMag-SELEX was performed as recommended by Stoltenburg *et al*. (2005) (22). Before each round, library or each ssDNA pool was heated at 90°C for 10 min, immediately cooled, and kept at 4°C for 15 min, then shortly incubated (5–8 min) at room temperature before their application in the binding reaction. To enhance the specificity, as counter selection, 500 μL of the B/W buffer containing 5.2 nmol of the random oligonucleotide library were incubated for 5.5 hours with a mix containing 50 µL of naked Ni-NTA beads and 50 µL of naked GSH magnetic beads without agitation. Then, the depleted library (around 4.8 nmol) was incubated with 200 µg of the target protein coated on beads at 22°C for 1 hour on a rotating wheel. The unbound oligonucleotides were removed by five washing steps with 500 μL B/W buffer. Then, the bound oligonucleotides were eluted by incubating the binding complex thrice with 200 μL elution buffer (40 mM Tris-HCl pH 8, 10 mM EDTA, 3.5 M urea, 0.02% Tween 20) at 90°C for 10 min.

The pooled oligonucleotides were recovered in 30 µL of water using Monarch® PCR & DNA Cleanup Kit (Oligonucleotide Cleanup protocol). All the selected ssDNA were amplified in 20 parallel PCR reactions. Each contained 1X of reaction buffer, 1X of stabilizer solution, 0.2 mM dNTPs each, 1 µM of each primer (with FAM and HEGL), 30 µL of the ssDNA pool recovered from the selection, and 1.25 U NeoProof DNA Polymerase in a volume of 50 µL. The amplification conditions were 5 min at 95°C and 30 cycles of 30 seconds at 94°C, 30 seconds at 47°C, 30 seconds at 72°C, then 5 min at 72°C after the last cycle (22). This resulted in the production of double-stranded DNA (dsDNA) fragments with a FAM modification at the 5′-end of the relevant strand and a poly-dA20 extension at the 5′-end of the complementary strand. To separate the relevant DNA strands from the double-stranded PCR products after the amplification step, the PCR product was purified as described above followed by 12% denaturing PAGE containing 7 M urea (**Supplementary Figure 1**). The fluorescein-labelled relevant DNA strands were identified in the gel by using the ChemiDoc™ imaging system and were cut out under a UV transilluminator. The gel was crushed and the DNA was eluted twice from the crushed gel with 200 µL of diffusion buffer (10 mM Tris, 25 mM sodium chloride, 1 mM EDTA, pH 8), at 70°C for 1 hour followed by 30 minutes, with mild shaking (1500 rpm). After recovering of the ssDNA as previously described, a new selected and fluorescein-labelled ssDNA pool was ready for the next selection round of the SELEX process.

After the first selection round, around 200 pmol from the previous round was used in the next round, starting with the binding reaction of the oligonucleotides to the target protein. The recovered ssDNA was quantified by absorbance at 260 nm using Nanodrop 2000. One negative selection (NS) and 12 positive selection rounds were performed to enrich target-specific oligonucleotides (aptamer candidates). The condition of each round is summarized in **Supplementary Table 2**.

### Cell-SELEX round of selection

The combination of Flu-Mag SELEX with Cell-SELEX was applied by using full bacteria instead of magnetic beads at the 10^th^ round of selection. *Borrelia* cells were grown in 50 mL Falcon™ conical tubes with a culture volume of 45 mL of Barbour-Stoenner-Kelly (BSK)-H medium at 34°C without agitation. After 15 days, 10 mL of the culture were centrifuged at 9 000 x g for 10 minutes and washed 3 times using the B/W buffer. After heating and cooling the ssDNA pool from the last round of selection, 200 pmol were incubated in 500 µL of the B/W buffer. 2 washing steps were performed with 500 µL of the B/W buffer including centrifugation at 9 000 g for 10 minutes. A heat elution was performed thrice in PBS at 95°C for 10 minutes. The bound ssDNA was recovered as previously described.

### Melting curve analysis

About 100 ng of PCR product from each SELEX cycle were purified with Monarch® PCR & DNA Cleanup Kit and subsequently incubated with 2× Neogreen qPCR Master Mix in a total volume of 20 µL. For melting curve analysis, the overall sample temperature was firstly increased at 95°C for 7 minutes, then the temperature was decreased at 40°C for 2 minutes followed by 5 minutes at 4°C. The sample temperature was increased at 40°C for 2 minutes, then increased gradually, from 40 °C to 95 °C at a step size 0.5 °C/2 seconds. Changes in sample fluorescence intensity were monitored continuously and melting peaks were smoothed and calculated with the cycler software.

### Beads and Dot Blot assays

The recognition of CspZ by the ssDNA pool from the different rounds of selection was characterized by two binding tests: beads assay and Dot Blot. For the beads assays, the fluorescein-labelled oligonucleotides from the library and from rounds 3, 7, 9 and 12, were heated to 95°C for 10 min and rapidly cooled to 4°C, then incubated at room temperature. Approximately 8 pmol of each pool were incubated with 12.5 µg of CspZ (i.e. 10 µL of coated beads) for 30 minutes at 22°C in a total volume of 100 μL of the B/W buffer. After the binding reaction, the unbound aptamers were removed by 5 washing steps with 500 µL of the B/W buffer. Heating elution at 95°C of bound oligonucleotides was then performed with 110 μL of SELEX elution buffer. 100 µL of the eluate were then transferred in 96-well black microtiter plates and the fluorescence was measured using VANTAstar. The FAM-labelled ssDNA library was used as a negative control.

For the Dot Blot technique, 0.25 nmol (i.e. 7 µg) of CspZ were coated on nitrocellulose membrane and dried 1 hour at 37°C. The membrane was blocked with 10% bovine serum albumin (BSA) in PBS buffer for 2 h at RT. Next, after 3 washings with PBS-Tween 0.1% and 2 washings with the B/W buffer for 5 minutes at RT, 8 pmol of ssDNA pool from the rounds of selection were added to each dot and incubated for 1 h at 22°C, then washed 3 times with the B/W buffer. The FAM-labelled ssDNA library was used as a negative control. For positive control, dots coated with 7 µg of His-tagged CspZ were incubated with anti-His antibody-FITC conjugate (1:500 diluted in PBS), then incubated for 1 hour at 37°C. The fluorescence signals were detected by the ChemiDoc™ imaging system.

### Next-generation sequencing (NGS)

NGS sequencing was outsourced by Paris Brain Institute, ICM. ssDNA pool from each round of selection have been previously amplified by PCR using the aforementioned primers and purified as previously described. Libraries were prepared according to standard protocols. The DNA library preparation was realized following the manufacturer’s recommendations (HyperPlus DNA kit from ROCHE). Final samples pooled libraries were sequenced on ILLUMINA Miseq with 2*150 micro cartridge (2x4 Millions of 150 bases reads), corresponding to 2 X 300000 reads per sample after demultiplexing. After recovering all the sequences from the NGS, an initial trimming has been proceeded using Cutadapt (34) by specifying the 5’ and 3’ primers, an error rate of 0.3 and removing the 0-length sequences. Additional filtration has been performed based on a length and quality analysis: after removing the primers, only the sequences with a length between 35 and 45 were retained. Moreover, given the high quality of the sequenced oligonucleotides, sequences showing a Q score < 30 on 3 or more independent positions were discarded. An in-house Python algorithm was employed to cluster identical sequences.

### Dot Blot, Biolayer Interferometry (BLI) assays and Surface Plasmon Resonance (SPR)

For the Dot Blot approach, 0.16 nmol (i.e. 4.5 µg) of CspZ were coated. Nitrocellulose membranes were blocked with BSA and washed. Then, 500 nM oligonucleotides conjugated with biotin were added to each blot. Revelation was performed using Biotin Polyclonal Antibody, HRP antibody (Bethyl Laboratories, A150-111P) 1/10 000 diluted. Similar conditions were used for FhbA and BSA (0, 16 nmol of coated proteins). Secondary antibodies used for positive controls revelation were His-probe (H-3) HRP antibody (SantaCruz, sc-8036-HRP) diluted 200-fold both for CspZ and FhbA. The chemiluminescence signals were detected by the ChemiDoc™ imaging system and quantified by integrated intensity using ImageJ software.

All BLI experiments were performed on an Octet HTX at the Robotein^®^ facility (part of Instruct ERIC) at 30°C using 384-well black polypropylene microplates (Greiner BioOne, Belgium), with a final volume of 80 μL per well. For all experiments, streptavidin (SA) biosensors (Sartorius: ref: 18-5019) were used for the immobilization of biotinylated oligonucleotides. The SA biosensors were hydrated in the B/W buffer for at least 10 min before use. A baseline step of 60 s was then performed in the B/W buffer. Next, biotinylated oligonucleotides were loaded onto the biosensors for 300 s using different oligonucleotides concentrations ranging from 0.125 to 1 µg/mL. A quenching step was performed by incubating the loaded biosensors in 10 μM biocytin for 300 s. A second baseline was then monitored for 120 s in the B/W buffer. The interaction with the purified CspZ was recorded for 600 s using different protein concentrations ranging from 0.26 µM to 50 µM depending of the considered aptamer (diluted in the B/W buffer supplemented with 0.1% BSA). The dissociation was recorded using the same buffer for 2 000 s which corresponds to an approximate period of time sufficient to dissociate 5% of formed complex (based on the determined k_off_ value). Recorded data were corrected by subtracting the signals from a reference sensor immobilized with oligonucleotides but without any analyte (B/W buffer-BSA). The binding of CspZ, FhbA and nanobody to SA biosensors without any aptamers was checked and do not show any binding (data not shown). All the data were analysed using the Octet Data Analysis HT software version 12.0, and fitted using a 1:1 interaction model with a global fit.

Regarding SPR assays, SA sensor chips were used in all the assays and conditioned using 1 M NaCl in 50 mM NaOH following the Biacore wizard. Subsequently, each aptamer was resuspended in the B/W buffer at a concentration of 1 µg/mL. The SA couplings were performed using the B/W buffer as the flow buffer throughout. Briefly, DNA aptamers were used as ligands by coupling via streptavidin/biotin procedure until targeted levels of 115 resonance units (RU) were reached. A control flow cell was similarly prepared but without injecting any protein. *Borrelia* recombinant proteins CspZ, FhbA (as functional mimic protein control), and BSA were used as analytes. Kinetic measurements were performed by injecting several amounts of *Borrelia* proteins, ranging from 0.25 µM to 15 µM at flow rate of 20 μL/min at 25 °C. Single cycle kinetics (SCK) were performed to avoid regeneration between injections due to concerns about aptamer denaturation with unknown refolding kinetics. Association time for each concentration was applied for 1000 seconds, followed by a dissociation time of 30 minutes before regeneration. Surfaces were regenerated by two injections of 60 μL of 100 mM NaOH. The blank run was subtracted from each sensorgram, prior to data processing using BIAcore T100 evaluation software. After the procedure, a logarithmic Langmuir 1:1 binding model and the simultaneous k_on_/k_off_ fitting of the BIAevaluation software was applied (Biacore, AB).

### Flow-cytometry and epifluorescence microscopy

To evaluate the ability of selected aptamers to bind available epitopes on bacteria, flow cytometry experiments were performed using a CytoFlex flow cytometer (Beckman Coulter Life Sciences, Indianapolis, USA) and the CytExpert software. After 8 days in BSK-H medium at 33°C, *Borrelia* culture was harvested by centrifugation at 8000 x g for 5 minutes and resuspended ten times in the starting volume of B/W buffer. After centrifugation, cells were resuspended in a volume corresponding to one twentieth of the culture volume in B/W buffer. 25 µL of this suspension (corresponding to 500 µL of initial culture) were added to 100 µL of polyclonal anti-CspZ antibody (Rockland, reference 200-401-C19, 1/200), and revealed by anti-rabbit antibody as secondary antibody (Invitrogen/Thermofisher, reference 31584, 1/750) for positive control. The secondary antibody alone was used as negative control. Binding of aptamers to *Borrelia* was tested using 25 µL of bacteria mixed with 100 µL of FAM-conjugated oligonucleotides, resulting in a final aptamer concentration of 500 nM. Whether the probe considered, after a 1-hour incubation at room temperature (RT) under static conditions, bacteria were washed with B/W buffer. Fluorescence of the samples was then measured using flow cytometry: Events (>150 000) corresponding to the cells were recorded for *B. burgdorferi* at a flow rate of 10 µL per minute for each treatment, and the emitted fluorescence resulting from FAM excitation at 488 nm was collected using a 525/40 bandpass filter. A threshold was applied for a SSC value greater than 1 000 (height), and gains were set as follows: FSC = 165, SSC = 400, and FITC = 240. Data were analyzed using the Cytexpert 2.0 software. Measurements were performed in triplicate.

Epifluorescence microscopy was performed on *Bb* B31 strain using the same preparation as applied for the flow cytometry experiments. Imaging was performed using an Olympus IX73 inverted microscope, and image analysis was conducted using CellSens Dimension software.

## RESULTS AND DISCUSSION

### Purification and proteins immobilization on magnetic beads

In this study, two recombinant CspZ were used for further selection of aptamers: His-tagged and GST-fused CspZ. Both recombinant proteins were expressed in *E. coli* BL21(DE3) and purified as previously optimized (31). SDS-PAGE analysis of the soluble extract and fractions after purification revealed a prominent band at approximately 48 kDa for CspZ/GST fusion protein and 28 kDa for His-tagged CspZ, as expected. Both proteins were purified at 88% and 99% (31) respectively as indicated by SDS-PAGE, Western blot and HPLC analysis (**Supplementary Figure 2**).

For protein immobilization, each protein was independently coated onto two distinct sets of magnetic beads in order to alternate the capture systems and avoid selection of oligonucleotides directed against the beads and the protein tags. Different amount of required purified proteins (400, 500, and 650 µg) were tested to ensure the saturation of magnetic beads. Unbound proteins were removed by washing and quantified as previously described. Results indicated that regardless of the initial amount of incubated CspZ, about 200 µg of protein were immobilized, suggesting complete saturation of the bead surfaces (**Supplementary Figure 3**). Therefore, 400 µg of CspZ was sufficient to saturate beads for the next steps.

### Nucleotide distribution and diversity of the initial/unselected DNA library

Compared to RNA oligonucleotides, ssDNA oligonucleotides exhibit greater stability for long-term storage, notably due to their inherent resistance to enzymatic degradation by nucleases. This enhanced stability makes ssDNA a more convenient molecule for subsequent applications. Therefore, the SELEX procedure was performed using a ssDNA oligonucleotides library. In this study, a randomized core region of 40 nucleotides (nt) was used with an equal content of all four nitrogenous bases, providing high sequence and structural diversity. Flanking regions consisted of 18 nucleotides and ensured the required amplification by PCR. The quality of the initial library was verified by NGS sequencing using an Illumina MiSeq sequencer. Nearly all oligonucleotides were at the expected length of 76 nucleotides (*i. e.* 40 nucleotides in the random region, **Supplementary Figure 4**). NGS analysis revealed a diversity of approximately 85%, defined as the total number of unique sequences over the number of total sequences. In addition, after clustering, the most abundant sequence exhibited a maximum of only four identical reads, further supporting the low effective redundancy which likely arises from PCR amplification prior to sequencing. Moreover, in accordance with our request to get library with equal content of each type of nucleotide, NGS analysis revealed a minor bias in the nucleotide distribution. Indeed, adenine constituted 19% of the random region, followed by cytosine at 20%, thymine at 28% and guanine at 33%. The detailed nucleotide composition for each selection round is provided in **Supplementary Table 3.** Bearing in mind that initial library composition can influence SELEX outcomes and favour specific structures such as stem-loops (35–37) or G-quadruplexes, the diversity of library was validated and the SELEX process to select aptamers against CspZ was thus performed.

### Aptamer selection against CspZ

Using the coated magnetic beads and the library described above, selection against the CspZ protein was performed using the cross-over-SELEX approach (22). Twelve rounds of selection were conducted in total (**Figure 1a**). Three rounds of selection were carried out against the 6His-CspZ protein using FluMag-SELEX strategy. Subsequently, in rounds 4 and 5, the oligonucleotides pool was incubated with GST-CspZ. By switching from Ni-NTA to GSH capture system, the aptamers binding to the tags (Histidine or GST) were removed (38), and the oligonucleotides were enriched for those specifically targeting the CspZ protein itself. Moreover, as the molecular weight of GST-fused CspZ is roughly twice that of His-tagged CspZ, the effective protein concentration for capture likely decreased by half when switching the tag system, given that a constant 200 µg of target protein immobilized per bead. Consequently, these rounds constitute a selection pressure. Following the established pattern in the first three rounds of selection, rounds 6 to 9 targeted the His-tagged CspZ. In round 8, the number of washes was increased from 5 to 8, and the incubation time was reduced from 1 hour to 30 minutes, introducing a new selection pressure. The tenth round was conducted directly on *Borrelia burgdorferi* B31 cells using Cell-SELEX, in order to achieve selective enrichment of oligonucleotides targeting CspZ epitopes available on the bacterial surface. Finally, rounds 11 and 12 were once again carried out against the His-tagged CspZ protein, with the goal of enhancing specificity. During each selection round, ssDNA bound to CspZ was eluted via salt and heat denaturation.

**Figure 1:**
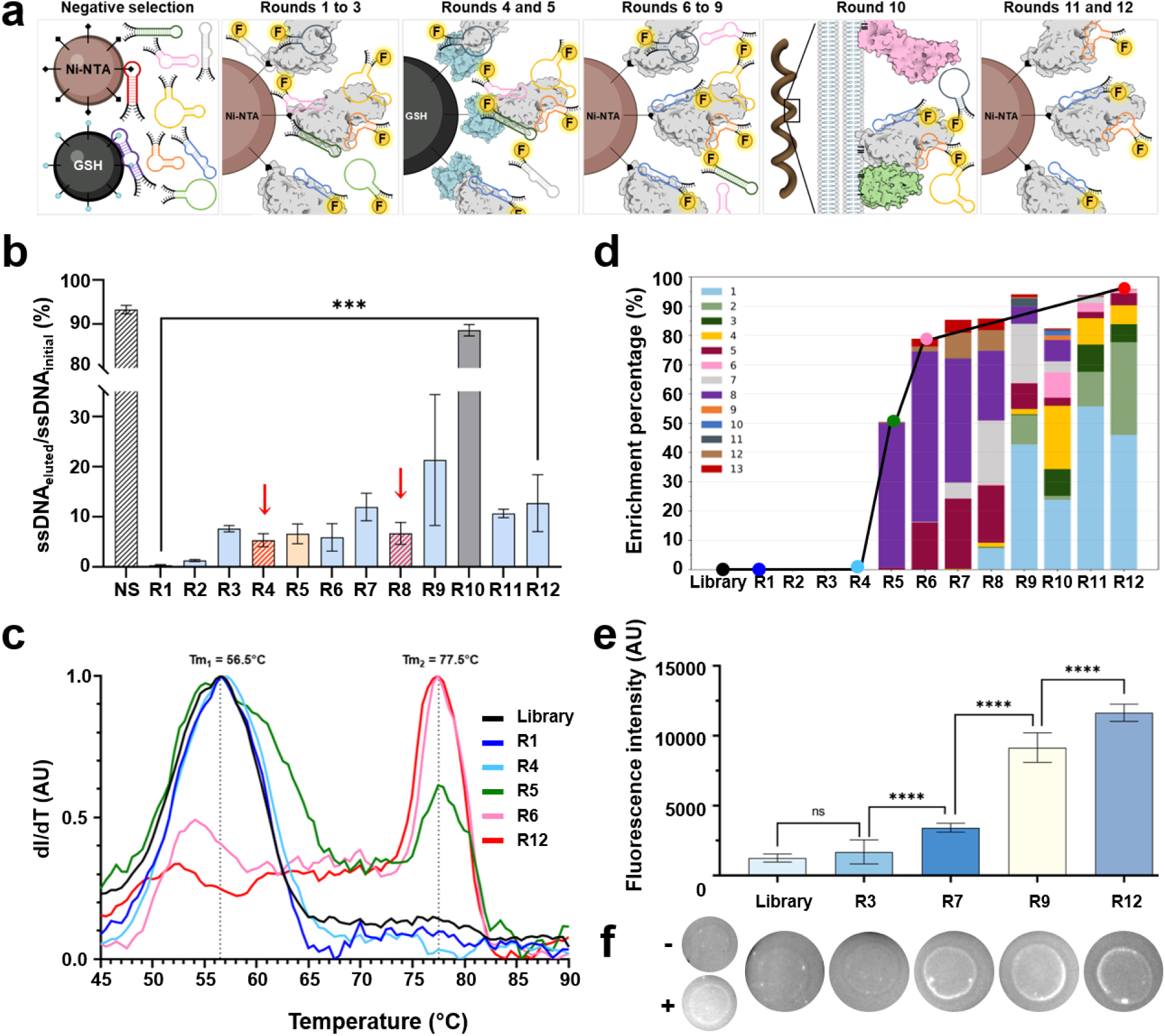
**a** – Schematic representation of the SELEX pipeline. **b** – Enrichment of CspZ-specific oligonucleotides during the cross-over-SELEX. In round 0, a negative selection (NS) has been performed (black hatched bar). Blue bars represent selection on Ni-NTA magnetic beads. Yellow bars represent selection on GSH magnetic beads. Round 10 with grey bar corresponds to a cell-SELEX round. The red arrows and hatched bars in red, represents pressure of selection. The error bars represent the standard deviation associated with multiple Nanodrop concentration measurements. The statistical analysis employed in this experiment was a two-sample t-test (with ***=p<0.001). **c** – Assessment of the SELEX process using melting curve analysis (first derivative dI/dT of fluorescence signal). Curves correspond to the measured Tm, after PCR, of dsDNA from the initial SELEX library and the dsDNA from selection rounds 1, 4, 5, 6 and 12. **d** – Identical sequences enrichment (continuous line) and clustering (histogram) as a function of the selection round. Only sequences/clusters with an occupancy > 1.5% (for at least one round of selection) are reported. **e** – Fluorescence intensity (AU) after beads assay for selected rounds of selection. Background fluorescence with the B/W buffer recovered after performing the same experiment with naked beads has been subtracted. The error bars represent the standard deviation associated with the mean fluorescence intensity from 3 replicate experiments. The statistical analysis employed in this experiment was a two-sample t-test (with ****=p<0.0001). **f** – Dot Blot analysis on selected rounds of selection. The negative control corresponds to the autofluorescence of His-tagged CspZ, while positive control corresponds to coated His-tagged CspZ revealed using anti-His antibody conjugated to FITC. For library or selection rounds tests, coated CspZ was revealed using each pool of oligonucleotides conjugated with FAM.

The absorbance of eluted ssDNA was monitored at 260 nm during the selection process. The gradual enrichment of oligonucleotides is illustrated in **Figure 1b**. The recovery percentage of bound ssDNA increased from 0.33% to 12.7% over the total duration of all rounds. This low recovery yield can be influenced by epitope concentration at beads’ surface. As expected, in rounds 4 and 8, where a selection pressure was applied, a decrease in bound oligonucleotides was observed, from 7.6% to 5.1% and 11.9% to 6.7%, respectively. For rounds 7, 9, 11, and 12, the amount of eluted ssDNA remained relatively consistent. In round 10, which was performed against the whole intact bacteria, a substantial amount (88.5%) of DNA was retained and eluted. Several potential explanations exist for this observation:

i. 10 mL of stationary-phase bacteria were collected and incubated with the oligonucleotides. In the stationary phase, the literature suggests that the bacterial density is around 1×10^8^ cells/mL (39), resulting in an estimated 1 x 10^9^ cells in this experiment. By using an equal number of beads and bacteria, the total surface area available for the bactera is significantly greater than that of the beads. This difference arises because the surface area of a typical spirochete (25 µm², assuming a cylindrical shape) is approximately eight times larger than that of a bead (3.1 µm²). Thus, a higher abundance of available target epitopes can be considered during the round using whole cells (round 10), leading to more bound oligonucleotides, contributing to the high eluted DNA yield.
ii. The presence of other proteins alongside the bacteria may have facilitated non-specific binding of oligonucleotides.
iii. The combination of heat and chemical agents employed for elution might have induced bacterial cell lysis, leading to the release of genomic DNA into the sample and finally to the overestimation of oligonucleotides dosage. Nevertheless, PCR amplification subsequently performed in SELEX procedure using specific primers, genomic DNA is expected to be eliminated. To confirm the loss of diversity during the selection process, both melting curve analysis and NGS sequencing were performed.

### Monitoring the SELEX procedure: melting curve and NGS analyses

The melting temperature (Tm) profiles of amplified dsDNA (from eluted ssDNA) from different SELEX rounds were evaluated to decipher the diversity evolution of aptamer pools, since melting curve analysis is a rapid, simple, informative, and efficient approach for assessing the convergence of oligonucleotide pool diversity. Indeed, this assay is based on the heat-induced denaturation of PCR products in the presence of an intercalating dye by measuring the sample fluorescence intensity. As previously described (40), the Tm of pools of oligonucleotides from successive selection rounds is expected to increase when the diversity decreases. As depicted in **Figure 1c**, the melting curves of the oligonucleotide library and amplified dsDNA from rounds 1 and 4 were characterized by one peak with a unique Tm (56.5°C), whereas for round 5, two broad peaks with different Tm (56.5°C and 77.5°C) were observed. Ultimately, for rounds 6 and 12 as the SELEX progressed, the shift of Tm is confirmed with a predominant peak at Tm 77.5°C.

In the random DNA library, the Tm is typically expected to reflect the average of Tm values of individual oligonucleotides. Each oligonucleotide contributes minimally to the overall diversity, resulting in a broad peak with a lower Tm. Similar results were observed with the first four rounds of selection. These broad peaks likely arise from the presence of less stable and imperfect heteroduplexes formed by incomplete hybridization within the flanking regions of the sequences. In contrast, the emergence of two distinct broad peaks in the melting curve of the fifth round indicates the initiation of enrichment. The higher Tm peak indicates the presence of more stable homoduplexes involving both the random core of oligonucleotides and their flanking regions. As the SELEX process successfully progresses, the formation of heteroduplexes is expected to diminish. Consequently, the gradual disappearance of the low Tm peak and the concomitant evolution of a single, distinct peak with a higher Tm reflect the success of SELEX approach and emergence of sequence clusters that potentially bind to CspZ (40–43).

To go further in the monitoring and the comprehension of the selection process, NGS sequencing from each round was performed. After trimming, quality control, and length filtering, NGS analyses provided a total of 74 824 to 141 575 sequences per selection round (**Supplementary Table 4**). The per round enrichment percentage was computed as the number of sequences with at least 1.5 % of redundancy compared to the total number of sequences. As depicted in **Figure 1d** (continuous line), an enrichment occurred abruptly at round 5, from approximately 0% to 50%, up to 80% at round 6. The enrichment is coherent with the results obtained from the melting curve analysis, where two distinct peaks were identified at round 5, corresponding to homo- and heteroduplexes. Taken together, whether through ssDNA elution assays, melting curve monitoring, or enrichment of sequences obtained by NGS, all results obtained are consistent with a successful selection of oligonucleotides binding to the target protein.

To reach a comprehensive representation of sequence redundancy, NGS allowed to characterize each oligonucleotide pool by identifying and clustering together identical sequences. Similar to the enrichment threshold applied, sequences with more than 1.5% redundancy for at least one round of selection have been clustered as shown in **Figure 1d**. The sequencing of the initial library validated that the most redundant sequence was detected at a maximum of four times out of a total of 105, 772 reads, *i.e.* less than 0.004%. From the round 5, a first cluster emerges, concomitantly with the oligonucleotide enrichment. Then, the cluster profiles evolve over the course of selection rounds and applied selection pressure. At the last round of selection, five clusters displayed high redundancy yields of 46% (Apta1), 32% (Apta2), 6.1% (Apta3), 6.5% (Apta4), and 4% (Apta5), collectively accounting for over 94% of the total sequences. All sequences are reported in **Supplementary Table 1**. Over the 12 selection rounds, 13 sequences were retained because they had an occurrence ≥ 1.5 %. In order to get preliminary information about the similarity of the retained sequences, a multiple sequence alignment using Clustal Omega (44) was performed. This tool performs a progressive multiple alignment by following a guide tree (**Figure 2a**) built on the base of the distances between sequences or group of sequences: close sequences are grouped and progressively aligned to farther sequences. The guide tree used by Clustal Omega to align the 13 sequences, together with the identity matrix (**Figure 2b**) generated from the alignment, highlighted a cluster of highly similar sequences, with identities exceeding 50%, i.e. Apta3, 4, 6, 9, 10, 11, and 12. Notably, aptamers 9 and 10 stand out with a high identity score of approximately 76%. Within this cluster, Apta3, 4, 6, 9, and 10 exhibited a high guanine nucleotide content (>50%) distributed in clusters of 2 to 4 guanines separated by 1 to 3 nucleotides, suggesting the potential formation of G-quadruplex structures.

**Figure 2:**
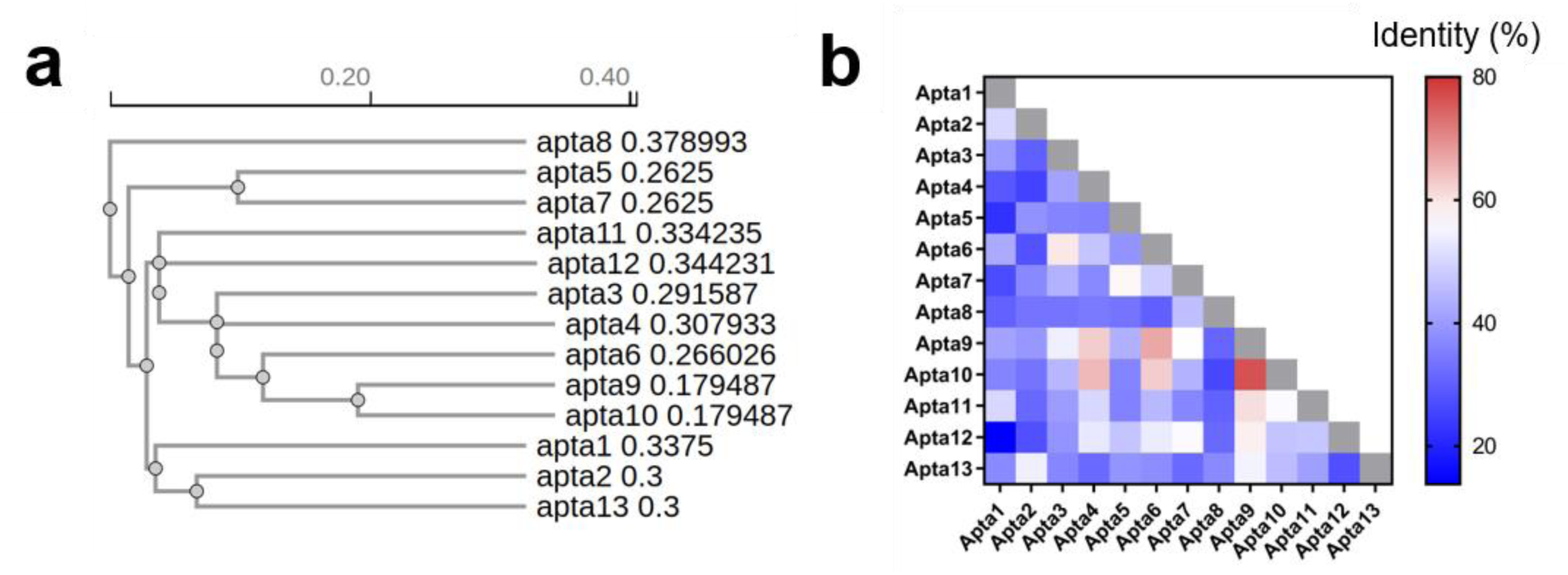
Data deduced from multiple sequence alignment of selected aptamers using Clustal Omega. **a** – Guide tree visualizing the relationships among sequences based on the alignment. **b** – Pairwise identity matrix depicting the percentage of identical residues between each pair of sequences highlighting the clusters of highly similar sequences in red gradient.

Beyond sequence enrichment, the functionality of oligonucleotides pools, *i.e.* those able to bind CspZ, has been assessed using both a beads assay and Dot Blot approach.

### Beads and Dot Blot assay for monitoring the enrichment of functional oligonucleotides targeting CspZ

Beads assays were performed on selected rounds of the selection procedure. As shown in **Figure 1e**, round 3 fluorescence intensity remained near background level. However, a significant increase in fluorescence signal was observed, ranging from 1 256 UA (library) to 11 644 UA (round 12), illustrating a successful enrichment in functional oligonucleotides.

Similarly, Dot Blot analysis was applied to fluorescent labelled ssDNA pools derived from SELEX rounds. Dot Blot analysis offers a rapid mean to assess oligonucleotide pool interaction with target proteins, and served as a preliminary screening tool (45–48). As shown in **Figure 1f**, initial library and round 3 samples display no fluorescence and are similar to the negative control. In contrast, fluorescence signal is detected on rounds 7, 9, and 12, mirroring the positive control. Dot Blot results clearly showed an enrichment of oligonucleotides able to bind to the target protein as the SELEX cycles proceeded.

### Biochemical and biophysical characterization of selected oligonucleotides

Thirteen candidate aptamers were identified (**Figure 1d**, histogram) for further characterization (numbered 1 to 13). For in-depth analysis, three distinct controls were considered: (1) a negative aptamer control (AptaCtrl), selected from the initial library but absent in the following SELEX rounds; (2) a negative protein control, BSA; (3) a second negative protein control which is an irrelevant nanobody containing a histidine tag. Additionally, FhbA, a functional mimic of CspZ, was considered as a specificity control. Produced by *Borrelia hermsii,* FhbA shares the same structural proportions (helix) and the same ability to bind FH and FHL-1 (31). As a pre-qualitative analysis, a Dot Blot has been performed with each individual candidate aptamer.

Dot Blot analysis with CspZ (**Figures 3a and 3d**) revealed interactions of varying intensities/integrated density for all oligonucleotides. As expected, the negative control AptaCtrl exhibited minimal signal intensity. Signals comparable to AptaCtrl were observed for oligonucleotides 1 and 2, despite those oligonucleotides were overrepresented in last round of selection. However, oligonucleotides 3 to 13 exhibited binding signals with CspZ. Oligonucleotides 3, 4, 6, 7, and 9 to 13 showed particularly strong signals, suggesting that these sequences may be predefined as interesting aptamers. When comparing with binding abilities to functional mimic FhbA (**Figures 3b and 3d**), only two oligonucleotides 3 and 11, exhibited strong binding signals, while faint signals were observed for all other aptamers (i.e. 1, 2, 4 to 10 and 12 to 13), suggesting weaker interactions. No interaction or very weak non-specific binding has been observed for all oligonucleotides regarding their interaction with BSA (**Figures 3c and 3d**).

**Figure 3:**
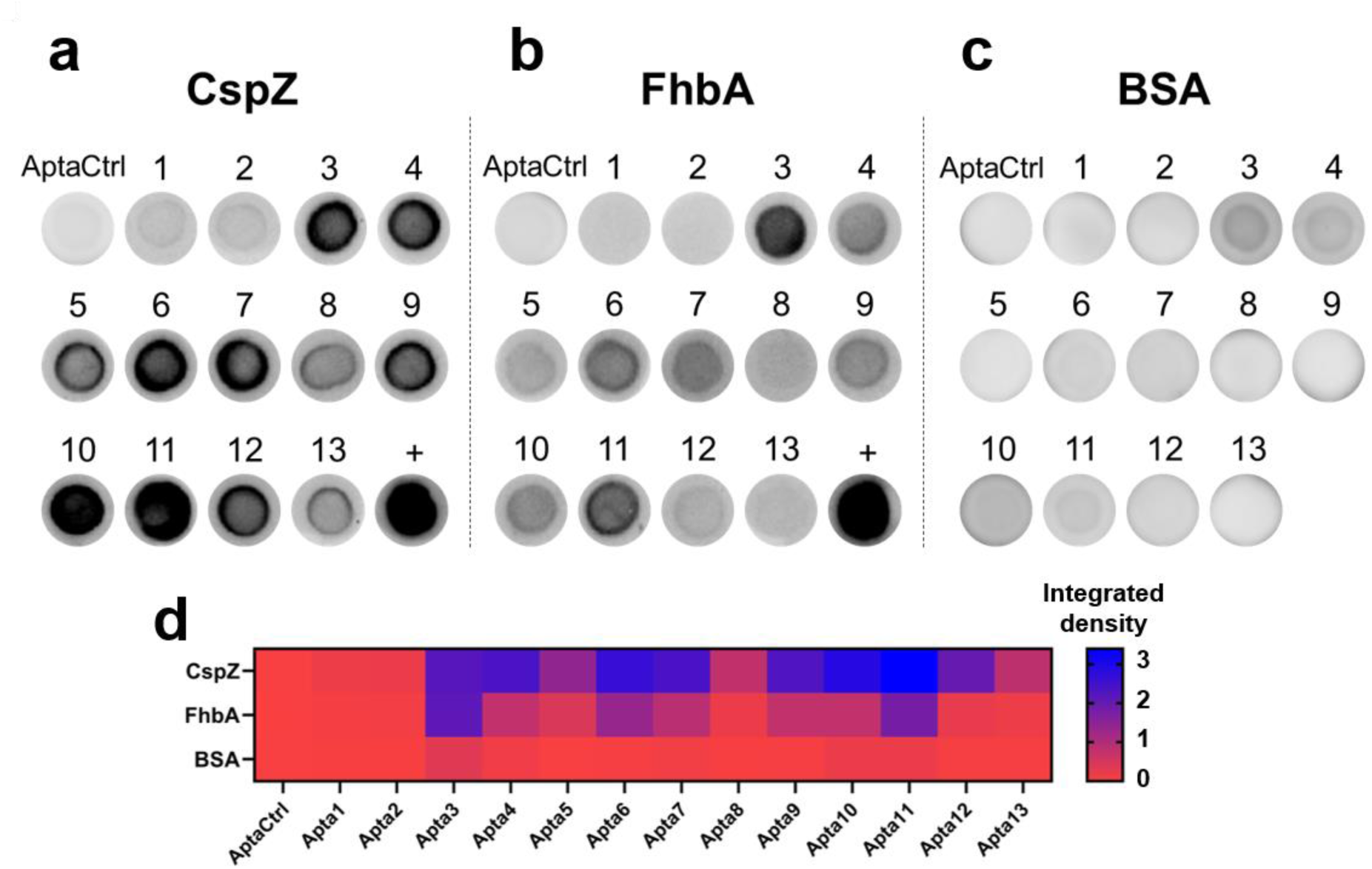
Dot Blot analysis for each selected oligonucleotide. The target protein was coated on membranes (i.e. **a** – CspZ, **b** – FhbA or **c** – BSA), and incubated with distinct biotinylated aptamers. Revelation was performed using anti-biotin polyclonal antibodies conjugated with HRP. Concentration of each partner remains constant in the various depositions. Positive controls (+) are revealed using specific antibodies conjugated to HRP.**d** – Integrated density of each dot with CspZ, FhbA and BSA, quantified using ImageJ software.

To confirm and further characterize the kinetics of interaction of oligonucleotides with CspZ, BLI was employed. Once again, AptaCtrl showed no interaction at 10 µM of CspZ or FhbA (**Supplementary Figures 5a and 5b**). Our results revealed binding of the selected candidates to CspZ, with stronger interactions observed for aptamers 3, 4, 6, 7, and 9 to 12 (**Supplementary Figure 5a)**. Binding was not observed for all the tested oligonucleotides with the His-tagged negative protein control (an irrelevant nanobody V_H_H), thus confirming the specificity of oligonucleotides to CspZ and not its histidine tag (**Supplementary Figure 5c**). Regarding specificity of these aptamers to FhbA, the corresponding BLI measurements were coherent with the Dot Blot data and showed some interactions with FhbA for aptamers 3, 4, 6, 7, 9, 10, 11 and 12 (**Supplementary Figure 5b**). To further characterize these observations, BLI measurements were also performed in more stringent conditions by altering the salinity of the buffer by tripling the NaCl concentration (**Supplementary Figure 6**). In these conditions, most interactions with CspZ were maintained (particularly for aptamers 3, 4, 10 and 11), while interactions with FhbA appeared to be lowered. It is worth noting that screening analysis does not allow to quantify the interaction (i.e. K_D_ constants). All together, these data suggest that it is possible to increase the specificity of the assay by increasing the ionic strength of the binding buffer. Beyond the screening, individual BLI experiments were conducted for each selected aptamer to determine the association/dissociation kinetic parameters enabling the calculation of the dissociation constant, as shown in **Table 1**.

**Table 1:** Quantitative analysis of the interaction between immobilized aptamers and CspZ by BLI.

| Sequence | K<sub>D</sub> (nM) | k<sub>on</sub> (M<sup>-1</sup>.s<sup>-1</sup>) | k<sub>off</sub> (s<sup>-1</sup>) | χ<sup>2</sup> | R<sup>2</sup> |
| --- | --- | --- | --- | --- | --- |
| **AptaCtrl** | No interaction observed at 10 μM CspZ |  |  |  |  |
| **1** | 565.4 ± 2.0 | 29.4 | 166.5 x 10<sup>-7</sup> | 117.4 | 0.999 |
| **2** | 534.5 ± 1.5 | 36.1 | 192.9 x 10<sup>-7</sup> | 169.3 | 0.999 |
| **3** | 47.4 ± 1.2 | 191.7 | 90.9 x 10<sup>-7</sup> | 14.9 | 0.997 |
| **4** | 37.8 ± 1.0 | 178.2 | 67.4 x 10<sup>-7</sup> | 16.4 | 0.998 |
| **5** | 1015.0 ± 1.6 | 34.4 | 349.1 x 10<sup>-7</sup> | 158.0 | 0.999 |
| **6** | 54.0 ± 0.3 | 153.0 | 82.7 x 10<sup>-7</sup> | 12.3 | 0.999 |
| **7** | 17.4 ± 0.3 | 246.1 | 42.8 x 10<sup>-7</sup> | 79.6 | 0.997 |
| **8** | 701.9 ± 1.9 | 33.7 | 236.2 x 10<sup>-7</sup> | 148.7 | 0.999 |
| **9** | 105.1 ± 0.7 | 244.4 | 256.8 x 10<sup>-7</sup> | 44.5 | 0.998 |
| **10** | 90.6 ± 0.8 | 324.3 | 293.7 x 10<sup>-7</sup> | 127.8 | 0.997 |
| **11** | 10.6 ± 0.1 | 843.8 | 89.3 x 10<sup>-7</sup> | 120.6 | 0.994 |
| **12** | 187.7 ± 0.6 | 157.1 | 294.8 x 10<sup>-7</sup> | 29.8 | 0.998 |
| **13** | 975.4 ± 1.5 | 33.8 | 329.5 x 10<sup>-7</sup> | 116.2 | 0.998 |

Associated sensorgrams are presented in **Supplementary Figure 7**. Both kinetic association and dissociation rates were found to be low. The equilibrium dissociation constants ranged from 10.6 nM to 1.0 µM.

The relatively high χ^2^ can be attributed to the unperfect match between the experimental data and the fits, especially at the start of the association curves. We hypothesized that some sorts of conformational changes in the aptamer could occur upon binding to CspZ, leading to these observed “biphasic” association curves. The binding affinities of selected aptamers with K_D_ deduced from BLI lower than 150 nM (i.e. aptamers 3, 4, 6, 7, 9, 10, and 11) for CspZ, were selected for further characterization by SPR. Single-cycle Kinetics (SCK™) was employed by injecting increasing concentrations of CspZ or FhbA (from 250 nM to 15 µM depending on the aptamer affinity). The sensorgrams presented in the **Supplementary Figure 8** and **Supplementary Figure 9**, respectively, and the kinetic parameters in **Table 2** demonstrate tight and stable binding of CspZ to the immobilized aptamers.

**Table 2:**
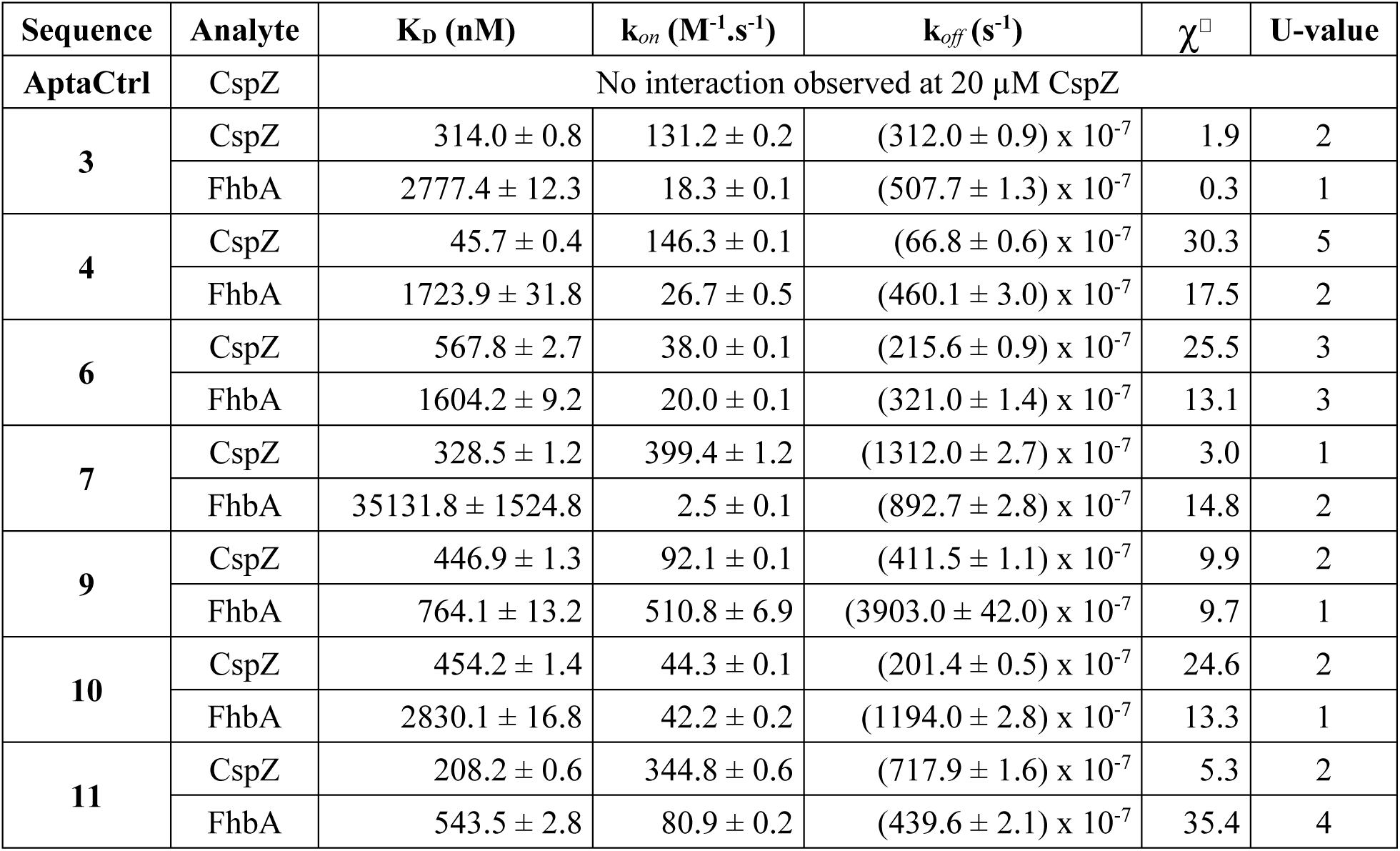
Quantitative analysis of the interaction between immobilized aptamers and CspZ or FhbA proteins by SPR.

| Sequence | Analyte | $K_D$ (nM) | $k_{on}$ ($M^{-1}.s^{-1}$) | $k_{off}$ ($s^{-1}$) | $\chi^2$ | U-value |
| --- | --- | --- | --- | --- | --- | --- |
| **AptaCtrl** | CspZ | No interaction observed at 20 μM CspZ | | | | |
| **3** | CspZ | $314.0 \pm 0.8$ | $131.2 \pm 0.2$ | $(312.0 \pm 0.9) \times 10^{-7}$ | 1.9 | 2 |
| | FhbA | $2777.4 \pm 12.3$ | $18.3 \pm 0.1$ | $(507.7 \pm 1.3) \times 10^{-7}$ | 0.3 | 1 |
| **4** | CspZ | $45.7 \pm 0.4$ | $146.3 \pm 0.1$ | $(66.8 \pm 0.6) \times 10^{-7}$ | 30.3 | 5 |
| | FhbA | $1723.9 \pm 31.8$ | $26.7 \pm 0.5$ | $(460.1 \pm 3.0) \times 10^{-7}$ | 17.5 | 2 |
| **6** | CspZ | $567.8 \pm 2.7$ | $38.0 \pm 0.1$ | $(215.6 \pm 0.9) \times 10^{-7}$ | 25.5 | 3 |
| | FhbA | $1604.2 \pm 9.2$ | $20.0 \pm 0.1$ | $(321.0 \pm 1.4) \times 10^{-7}$ | 13.1 | 3 |
| **7** | CspZ | $328.5 \pm 1.2$ | $399.4 \pm 1.2$ | $(1312.0 \pm 2.7) \times 10^{-7}$ | 3.0 | 1 |
| | FhbA | $35131.8 \pm 1524.8$ | $2.5 \pm 0.1$ | $(892.7 \pm 2.8) \times 10^{-7}$ | 14.8 | 2 |
| **9** | CspZ | $446.9 \pm 1.3$ | $92.1 \pm 0.1$ | $(411.5 \pm 1.1) \times 10^{-7}$ | 9.9 | 2 |
| | FhbA | $764.1 \pm 13.2$ | $510.8 \pm 6.9$ | $(3903.0 \pm 42.0) \times 10^{-7}$ | 9.7 | 1 |
| **10** | CspZ | $454.2 \pm 1.4$ | $44.3 \pm 0.1$ | $(201.4 \pm 0.5) \times 10^{-7}$ | 24.6 | 2 |
| | FhbA | $2830.1 \pm 16.8$ | $42.2 \pm 0.2$ | $(1194.0 \pm 2.8) \times 10^{-7}$ | 13.3 | 1 |
| **11** | CspZ | $208.2 \pm 0.6$ | $344.8 \pm 0.6$ | $(717.9 \pm 1.6) \times 10^{-7}$ | 5.3 | 2 |
| | FhbA | $543.5 \pm 2.8$ | $80.9 \pm 0.2$ | $(439.6 \pm 2.1) \times 10^{-7}$ | 35.4 | 4 |

As expected, AptaCtrl exhibited no detectable binding to either BSA, CspZ or its functional mimic FhbA. Furthermore, no interaction between Apta10 (arbitrarily chosen) and BSA (used as a control protein) was observed up to 20 µM BSA concentration (data not shown). Conversely, interaction of selected aptamers with CspZ was confirmed. When compared to BLI experiments, data are within the same range, though the ranking changes. Using the BLI approach, the most affine aptamer was Apta11 with a K_D_ of 10.6 ± 0.1 nM, followed by aptamers 7, 4, 3, 6, 10 and finally 9 with a K_D_ of 105.1 ± 0.7 nM. By SPR, Apta4 displayed the lowest K_D_ of 45.7±0.4 nM, followed by aptamers 11, 3, 7, 9, 10 and finally 6 with a K_D_ of 567.8 ± 2.7 nM. Experimental conditions, static solution *versus* continuous flow approach, and buffer compositions might explain such differences. Moreover, K_D_ obtained by SPR were generally higher than those from BLI assays. This discrepancy could be attributed to the flow-induced enhancement of dissociation rate during SPR experiments.

The analysis performed with the functional mimic FhbA suggest a weaker binding of aptamers as compared to CspZ, in the range of µM, characterized by lower association rate (particularly for aptamers 3, 7, and 10) and higher dissociation rate. Nevertheless, since binding was observed, these results suggest that it is possible to adapt the selection conditions to identify more specific aptamers, for example by including more stringent conditions such as increased salt concentration (see BLI data in **Supplementary Figure 6**). Finally, it is also worth to mention that the kinetic rates of binding are fairly low (≈10^2^ - 10^3^ M^-1^ s^-1^). This result is may due to the SELEX experimental conditions. Therefore, to improve the observed low k*_on_* rate, the SELEX process could be optimized by reducing the incubation time between CspZ and the oligonucleotide pools.

### Flow-cytometry and epifluorescence microscopy as a tool for recognizing *Borrelia*

Since the ultimate goal of the selected aptamers is to detect CspZ in its native conformation on the bacterial surface, *in vitro* experiments using bacterial cultures are necessary. Nevertheless, according to the literature, the CspZ protein is very poorly expressed *in vitro* (27, 49). Therefore, the expression profile of the gene encoding for CspZ during the growth of *Bb* strain B31 using RT-qPCR was determined to identify an optimal growth phase for further binding experiments. Although the results confirm that *cspZ* is weakly expressed during *in vitro* growth, RT-qPCR analysis revealed that an 8-day culture period is optimal for binding experiments, coinciding with the peak expression of *cspZ* gene (**Supplementary Figure 10**).

For further applications in diagnostics, not only aptamers have to be specific for their target, but also be able to recognize the target as it is exposed on the cell surface. To assess aptamers recognition of bacteria, flow cytometry analysis was employed. Polyclonal antibodies directed against CspZ, revealed by secondary antibodies, were used as positive control and confirmed the expression (at a low level) of CspZ on the bacterial surface (**Supplementary Figure 11)**. Each aptamer was independently tested (**Figure 4**).

**Figure 4:**
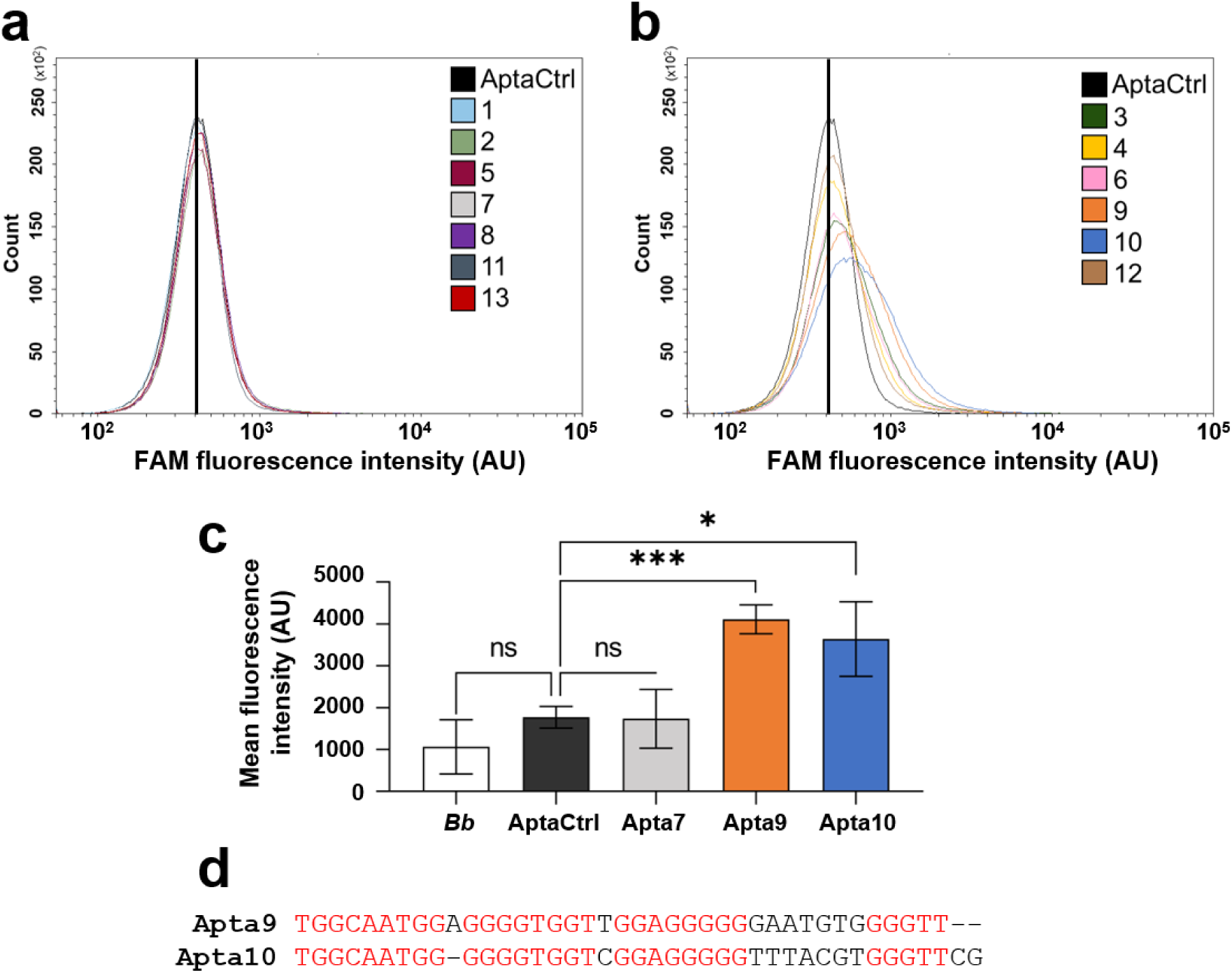
Recognition of *Bb*ss by selected aptamers using flow cytometry. Incubation with 500 nM of AptaCtrl in black **a** – Aptamers considered as not interacting with the bacteria *Bb*ss, mirroring the AptaCtrl. **b** – Aptamers considered as interacting with *Bb*ss in comparison with AptaCtrl. The black line highlights the negative control signal **c**– Recognition of *Bb*ss by selected aptamers (mean fluorescence intensity evaluated using flow cytometry). Incubation without aptamers (in white), or with 500 nM of AptaCtrl (black), Apta7 (grey), Apta9 (orange), or Apta10 (blue). The error bars represent the standard deviation associated with the mean fluorescence intensity from 3 independent experiments. Two-sample t-test was applied for the statistical analysis (*= p<0.05, and ***=p<0.001). d – Sequence alignment of Apta9 and Apta10, generated using Multalin (50). Identical nucleotides between the two aptamers are highlighted in red.

As illustrated in **Figure 4b**, six aptamers, namely aptamers 3, 4, 6, 9, 10, and 12 induced a shift in the fluorescence range. Aptamers 9 and 10 exhibit the most pronounced shift, in the same range of the antibody. In contrast, aptamers 1, 2, 5, 7, 8, 11 and 13 completely superimposed with AptaCtrl, suggesting that the epitope these aptamers target may not be accessible on the bacterial surface (**Figure 4a**). It is noteworthy that Apta7 and 11, that bind to FhbA similarly to Apta9 and 10 in dot blot experiments (**Figure 3b**), do not exhibit any interaction with the available epitopes of CspZ unlike Apta9 and 10. Indirectly, these results suggest that aptamers 7 and 11 also lack binding to other bacterial surface proteins. Consequently, the data gathered earlier using the functional mimic FhbA, which potentially indicated a non-specific interaction between aptamers and others borrelial surface proteins, could be considered as negligible.

A quantitative analysis of fluorescence signal deduced from flow cytometry experiments was then performed. Apta7, which binds to recombinant CspZ but not to whole bacteria (**Table 1** *versus* **Figure 4a**), served as an additional negative control. The mean fluorescence intensity of free *Borrelia* and FAM-Apta labelled *Borrelia* (with AptaCtrl, Apta7, 9, and 10) is presented in **Figure 4c**. Apta9 and Apta10 exhibited significantly higher fluorescence mean (4112 ± 348 AU and 3641 ± 889 AU, respectively) compared to AptaCtrl (1171 ± 257 AU). In contrast, Apta7 didn’t show any significant difference (1736 ± 699 AU). These findings corroborate the initial screening analysis, where Apta9 and Apta10 (**Figure 4b**) demonstrated binding to *Borrelia*, whereas Apta7 did not (**Figure 4a**). Remarkably, the alignment of Apta9 and 10 sequences (**Figure 4d**), that were indicated in the multiple alignment as the closest sequences (distance = 0.18, **Figure 2a**) and having the highest degree of identity (i.e. 76.32 %, **Figure 2b**), highlighted the redundance of guanine nucleotides with a specific distribution which may explain their comparable binding capacity to *Borrelia*.

To illustrate and validate the flow cytometry data, epifluorescence microscopy was performed on *Bb* (**Figure 5**). The microscopy data confirmed the ability of Apta9 and Apta10 to interact with targeted protein at bacterial surface in a low fluorescence intensity, as expected from flow cytometry data. Notably, the observed fluorescence distribution was similar to the recognition profile obtained with anti-CspZ antibody (30), suggesting that aptamers exhibit comparable binding affinity to the bacterial surface. In accordance with expectations, no signal was detected with AptaCtrl and Apta7.

**Figure 5:**
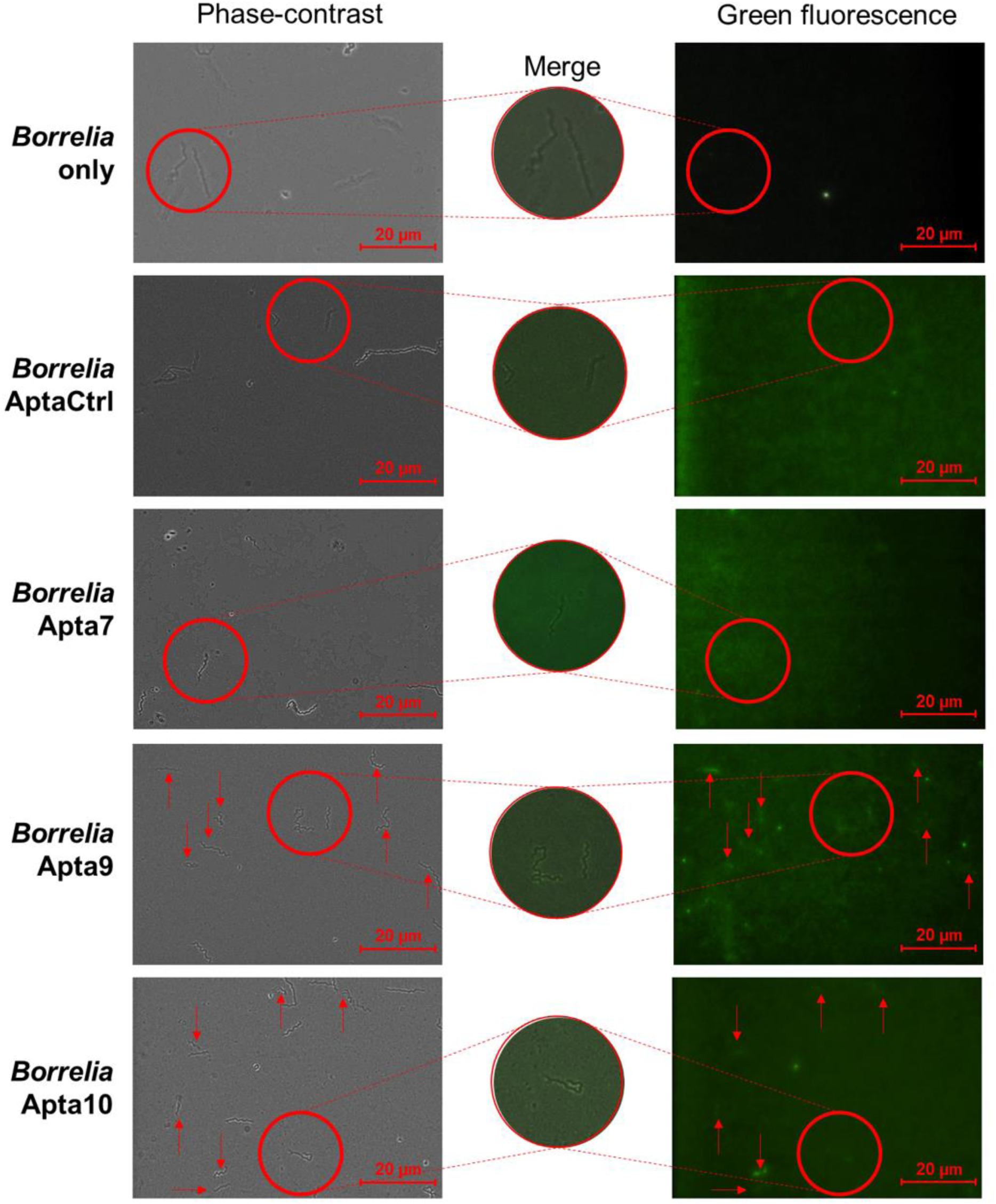
Epifluorescence microscopy of *Bb* bacteria (40x magnification). Images were acquired without aptamers (control) and using AptaCtrl, Apta7, Apta9, and Apta10, all conjugated with FAM. The red circle highlights the merged image (phase contrast and fluorescence) in the center. Red arrows point to some *Borrelia* bacteria exhibiting a fluorescence signal.

In this study, SELEX was employed and thirteen aptamers targeting the CspZ protein were successfully identified. While Apta1 and Apta2 were ultimately selected with high enrichment, it is noteworthy that their binding affinity is low. However, it is widely recognized that aptamer enrichment is not always directly correlated with binding affinity due to factors such as PCR bias, selection pressure, and aptamer folding and structure (33). Among these, seven aptamers exhibited high affinity (tens to hundreds of nanomolar range) towards CspZ, as determined by various assay formats (Dot-Blot, SPR, BLI).

Although this cross-over SELEX employed only a single round of cell-SELEX, additional cell-SELEX rounds could potentially improve aptamer specificity. However, the single-round approach was chosen to mitigate the risk of aptamers binding to other proteins on the bacterial surface, particularly in the absence of a knockout *Borrelia* strain. By applying this approach, two aptamers (Apta9 and Apta10), which also showed significant sequence identity, displayed specific binding not only to the soluble recombinant CspZ protein, but also to the native protein expressed at the bacterial surface. Data collected suggest that the specificity of the interaction can be influenced by several parameters. In particular, these include: (i) the differing expression environments of proteins *in vitro (*at different stages of the culture or with addition of blood for example) and *in vivo* (inside the tick, or in the host, at the beginning or the late infection), and (ii) the composition of the binding buffer such as the presence of detergents or salts. These findings establish a foundation of a proof of concept for the development of diagnostic probes with high affinity and specificity for CspZ, but also for all proteins displaying an available epitope. The direct detection of *Borrelia* antigens or whole bacterial cells holds promise for the early identification of LB (51). To our knowledge, only one study conducted by Tabb and colleagues, has investigated the detection of a *Borrelia* antigen using aptamers specifically targeting the OspA protein. Interestingly, the study demonstrated an aptamer with a nanomolar dissociation constant as determined by fluorescence anisotropy. Nevertheless, the recognition of surface-exposed epitopes of the bacterium by flow cytometry is not detailed. Both this study and the work of Tabb and collaborators highlights the potential of aptamers as LB diagnostic tools (8).Collectively, these results provide a strong proof-of-concept for developing novel diagnostic approaches. Indeed, aptamers, with their high target specificity and affinity, emerge as a promising alternative to antibodies. Their advantages include simple *in vitro* selection and production, ease of modification and conjugation, high stability, and low immunogenicity (14). In recent years, aptamer-based sensing platforms, known as aptasensors and relied on electrochemical, optical, or mass-sensitive principle, have been extensively investigated for diagnostic applications due to their significant advantages over traditional analytical techniques (52). Thus, the aptamer-based strategy presented here has the potential for broad applicability in the development of diagnostic tools for other bacterial biomarkers related to LB, aiming for example the detection of coinfections.

## DATA AVAILABILITY

All fastq files of SELEX are available in the Recherche Data Gouv website at https://recherche.data.gouv.fr/fr, DOI: 10-57745/RABFSQ.

## SUPPLEMENTARY DATA

Supplementary Data are available online.

## AUTHOR CONTRIBUTIONS

Mickaël Guérin: Conceptualization, Formal analysis, Investigation, Methodology, Validation, Visualization, Writing – original draft, Writing – review & editing. Marylène Vandevenne: Formal analysis, Methodology, Writing – review & editing. André Matagne: Formal analysis, Methodology, Writing – review & editing. Willy Aucher: Formal analysis, Methodology, Resources, Writing – review & editing. Julien Verdon: Formal analysis, Methodology, Resources, Writing – review & editing. Emmeline Paoli: Formal analysis, Methodology, Writing – review & editing. Jules Ducrotoy: Methodology. Stéphane Octave: Methodology, Resources, Writing – review & editing. Bérangère Avalle: Methodology, Visualization, Writing – review & editing. Irene Maffucci: Formal analysis, Methodology, Data curation, Visualization, Writing – review & editing. Séverine Padiolleau-Lefèvre: Conceptualization, Formal analysis, Funding acquisition, Investigation, Methodology, Project administration, Supervision, Validation, Visualization, Writing – review & editing.

All authors contributed to the article and approved the submitted version.

## ACKNOWLEDGEMENTS

The authors thank Prof. Marconi (USA), Prof. Kraiczy (Germany), Dr Brangulis (Latvia), and Dr Kogan (Finland) for providing the plasmids necessary for this study. The authors also thank Adeline Sellier for her technical assistance in protein production, and Dr Frédéric Ducongé for his valuable advice on SELEX analysis. We acknowledge the Robotein® platform of the BE Instruct-ERIC Centre for providing access to their equipment. NGS sequencing benefited from equipment and services from the iGenSeq core facility (Genotyping and sequencing), at ICM (Institut du Cerveau de Paris).

