## Supplementary for "Selection and characterization of DNA aptamers targeting the surface borrelial protein CspZ with high-throughput cross-over SELEX"

### Transcription level evaluation *cspZ* by RT-qPCR protocol

Frozen stock vials containing low-passage *Bb* strain B31 were thawed and inoculated into 2.5 mL BSK-H medium (Sigma) and incubated at 33°C until reaching a cell density of approximately $4x10^7$ spirochetes/mL. Then, spirochetes were cultured in BSK-H medium (13 mL into 15 mL Falcon™ tubes) at a starting concentration of $5.5x10^5$ spirochetes/mL. Daily monitoring of cell density was performed by spirochetes counting using dark-field microscopy. At different time (days 3, 5, 8 and 14), cells were harvested by centrifugation at 8,000 × g for 10 minutes and bacterial pellets were stored at -80°C prior to RNA extractions.

Total RNA was extracted from frozen pellets using the Direct-zol™ RNA MicroPrep kit (R2062; Zymo Research) according to the manufacturer's instructions. As recommended, a DNase I treatment was performed to remove contaminating DNA. The quality RNA integrity number ([RIN] > 7) and quantity of RNA were determined using the Agilent 4150 Tapestation system (Agilent Technologies). A total of 200 ng of RNA was standardized across samples from each experiment and converted to complementary DNA (cDNA) using the GoScript™ reverse transcription system (Promega) according to manufacturer's instructions. In this study, the *cspZ* gene forward and reverse primers were respectively 5'-GCCTACATTGAGAGTTTTGA-3' and 5'-GCCCCCTCAAGTTCTACAGC-3'. For the housekeeping gene, *flaB,* the forward and reverse primers were respectively 5'-TTGGAGCAAACCAAGATGAAGC–3' and 5'-TGCTGGCTGTTGAGCTCCTT-3'. qPCR was carried out using Takyon™ No Rox SYBR® MasterMix dTTP Blue (Eurogentec) with Forward and Reverse primers and a LightCycler 480 thermocycler. Primer sets for target genes were designed using the NCBI primer tool (https://www.ncbi.nlm.nih.gov/tools/primer-blast/). The amplification protocol comprised an initial denaturation step at 95°C for 5 minutes, followed by 45 cycles of amplification (denaturation 10 s at 95°C, annealing 10 s at 60°C, and extension 10 s at 72°C). Each run was performed with technical duplicates and melting curves were generated for each run to validate the production of specific amplicons. The data were analyzed using the ΔΔCt method, with normalization to *flaB* as a reference gene [1].

[1] T. Bykowski, M.E. Woodman, A.E. Cooley, C.A. Brissette, V. Brade, R. Wallich, P. Kraiczy, B. Stevenson, Coordinated Expression of Borrelia burgdorferi Complement Regulator-Acquiring Surface Proteins during the Lyme Disease Spirochete's Mammal-Tick Infection Cycle, Infect Immun 75 (2007) 4227–4236. https://doi.org/10.1128/IAI.00604-07.

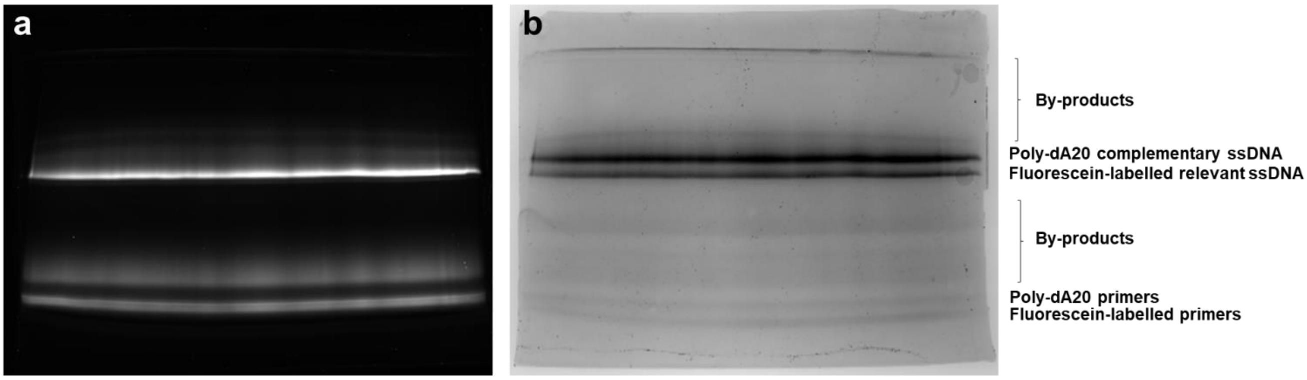


**Supplementary Figure 1 : 12% denaturing PAGE - 7 M urea. a – Fluorescence revelation. b – Methylene blue 0.02% staining.**

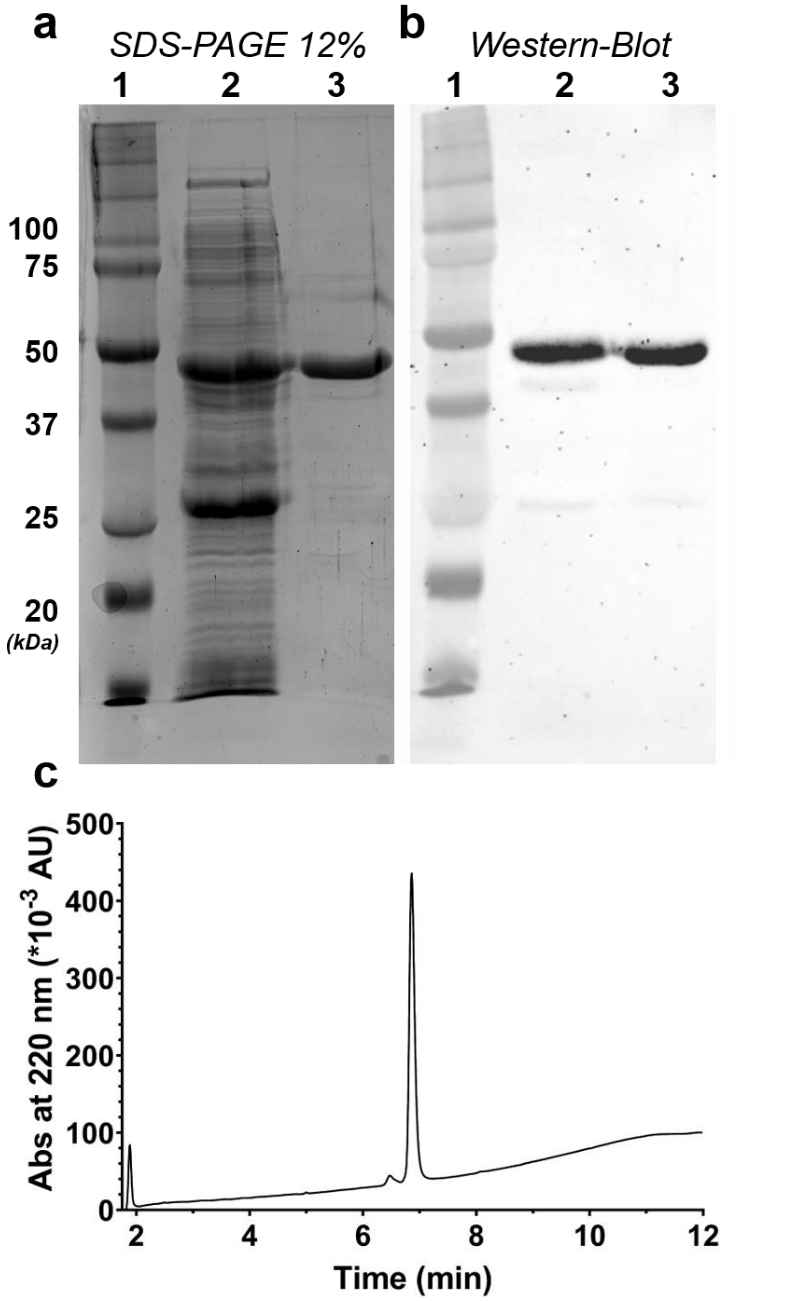


**Supplementary Figure 2 : Analysis of CspZ fused with GST. a – 12% SDS-PAGE gel stained with Coomassie Blue. Molecular weight marker (Precision Plus Protein Dual Color, #1610374, Biorad) (lane 1), migration patterns of soluble crude extract (lane 2) and purified CspZ protein (lane 3). b – Western Blot detected using horseradish peroxidase (HRP)-conjugated anti-GST tag antibody 1/1000 diluted (#sc-138, Santa Cruz). c – Purity evaluation of GST-fused CspZ by HPLC at 220 nm.**

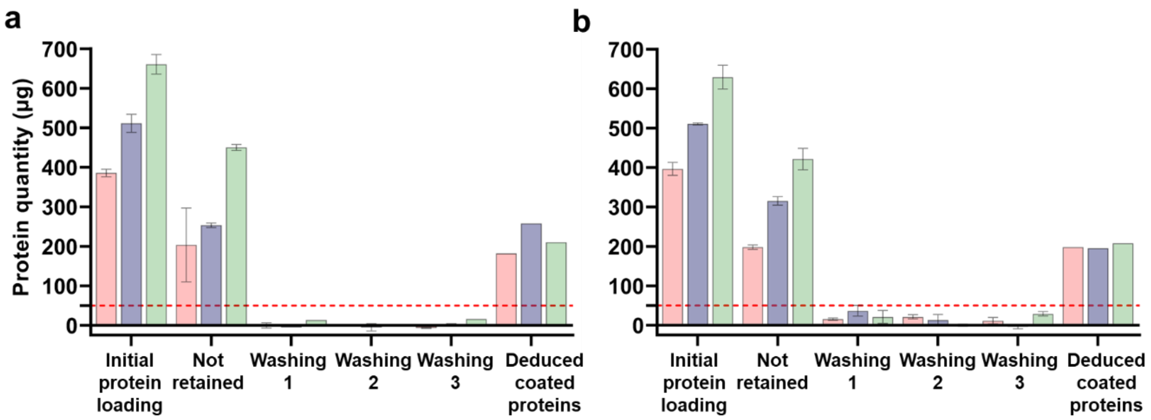


**Supplementary Figure 3 : Saturation conditions of magnetic beads by target proteins. a – Quantification of His-Tagged CspZ incubated with Ni-NTA beads, measured by BCA assay. In red, purple and green, loading of 390, 510 and 660 µg of CspZ respectively. b – Quantification of GST-fused CspZ incubated with GSH beads, measured by Pierce 660 nm. In red, purple and green, loading of 400, 500 and 630 µg of CspZ respectively. In both graphs, "deduced coated proteins" represents the difference between total proteins incubated with beads and unretained proteins. The red dotted line represents the limit of detection of the assays.**

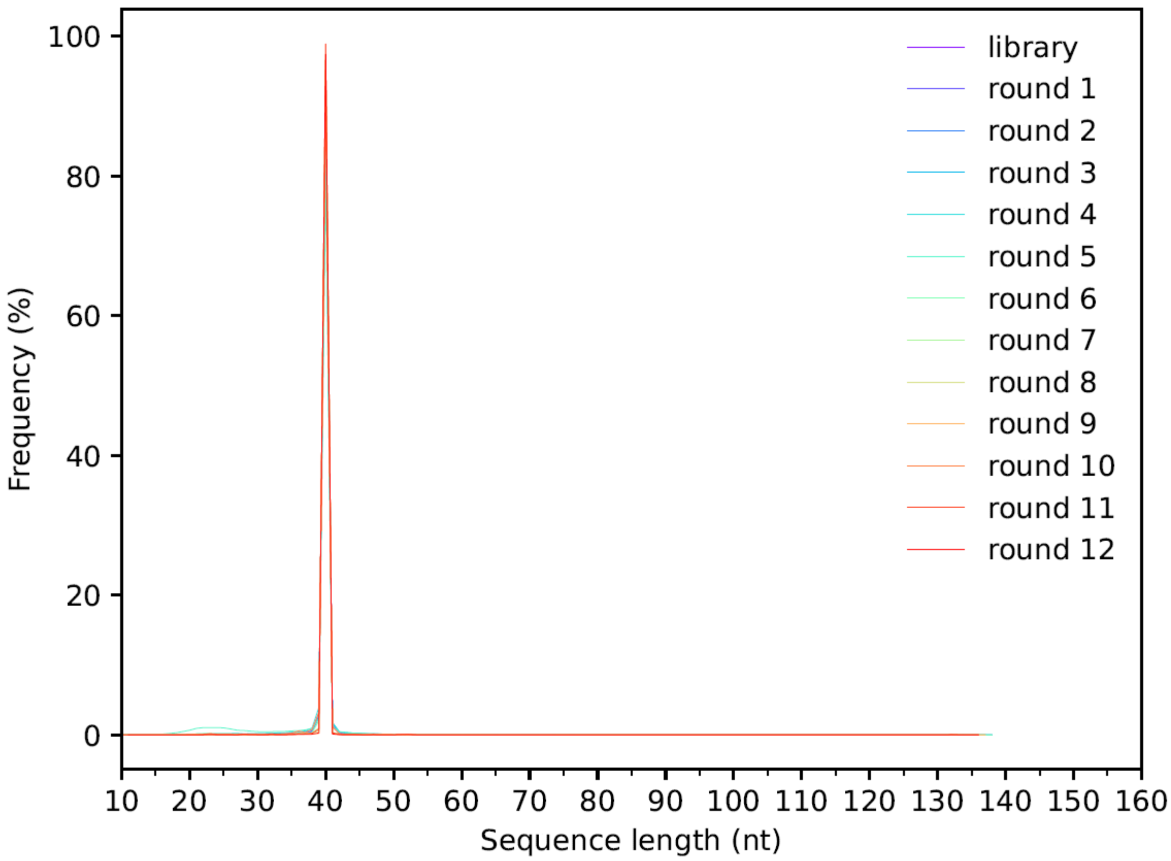


**Supplementary Figure 4 : Sequence length frequencies after the initial trimming.**

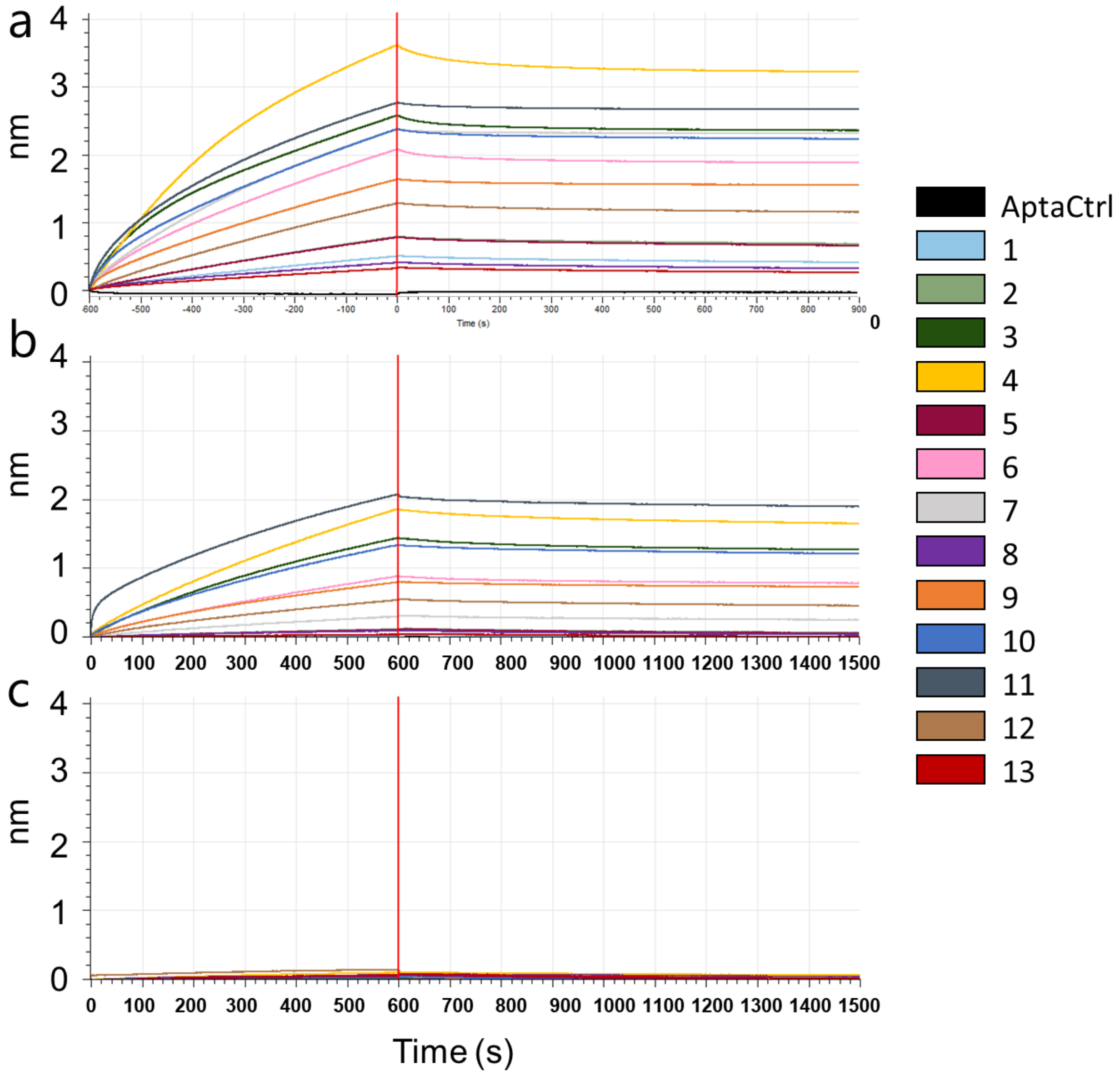


**Supplementary Figure 5 : Binding measurements performed by BLI showing the screening of all immobilized oligonucleotides (1 µg/mL) with different proteins. Association and dissociation times are 600 seconds and 900 seconds respectively. 10µM of the 3 tested proteins in 100mM NaCl binding buffer were used as analytes, i.e. (a) CspZ, (b) FhbA, and (c) an irrelevant nanobody.**

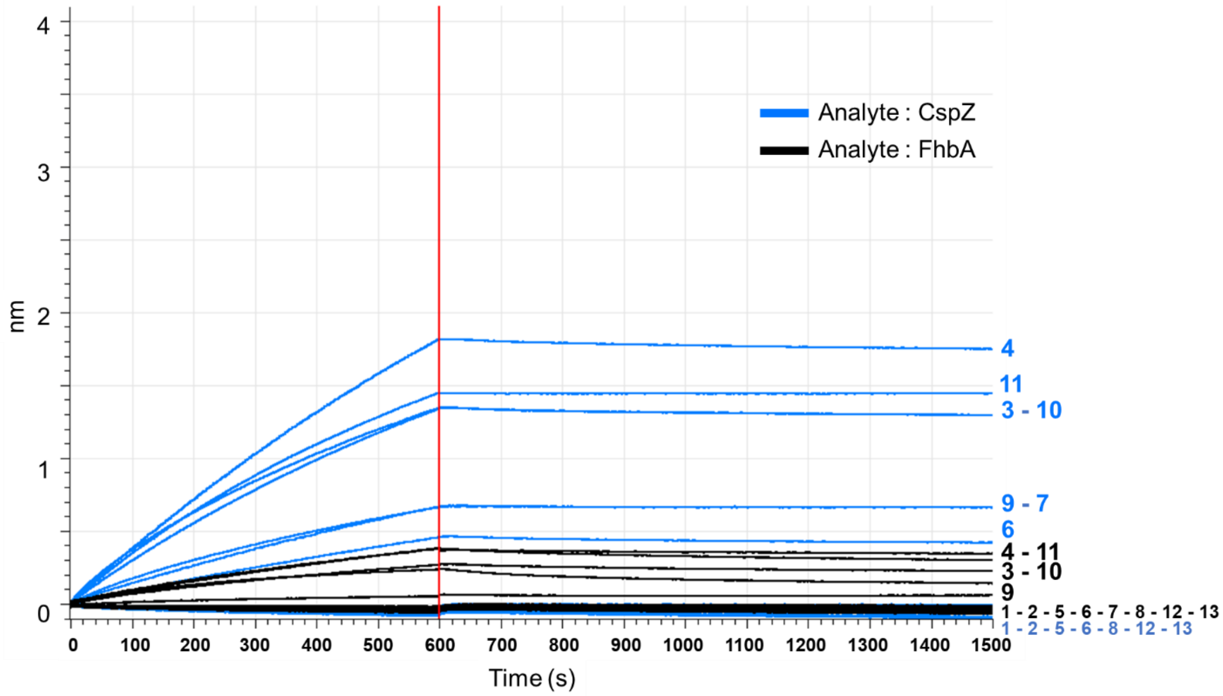


**Supplementary Figure 6 : Binding measurements performed by BLI showing the screening of all oligonucleotides (immobilisation level: 1 µg/mL) with 10 µM CspZ and 10 µM FhbA (in blue and black respectively) in 300 mM NaCl binding buffer. Association and dissociation times are 600 seconds and 900 seconds respectively.**

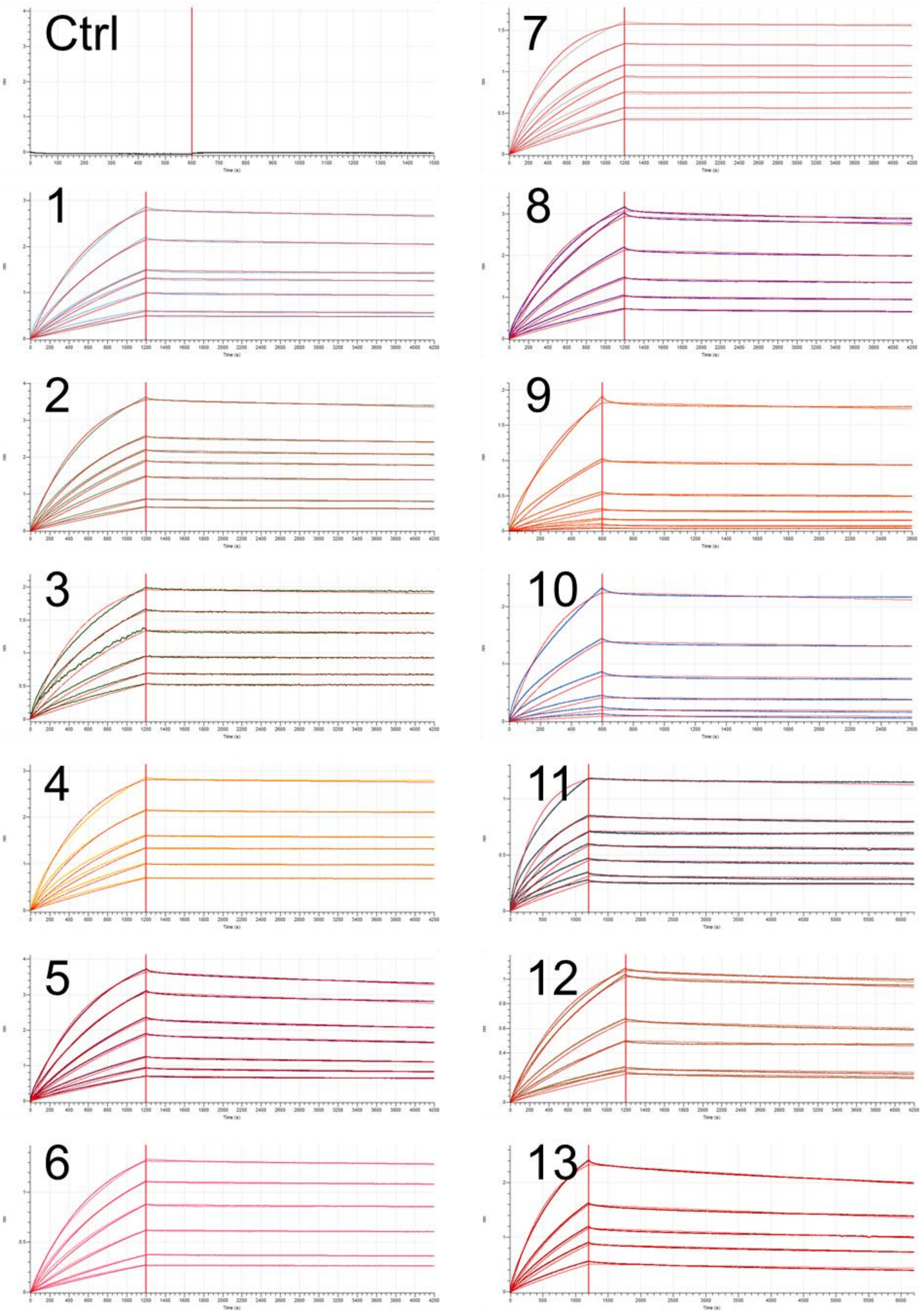


**Supplementary Figure 7 : BLI experiments for aptamers 1 to 13 with concentration of CspZ ranging from 0.26 µM to 50 µM depending of the aptamers. Red curves represent the fitted 1:1 model superimposed on the experimental data.**

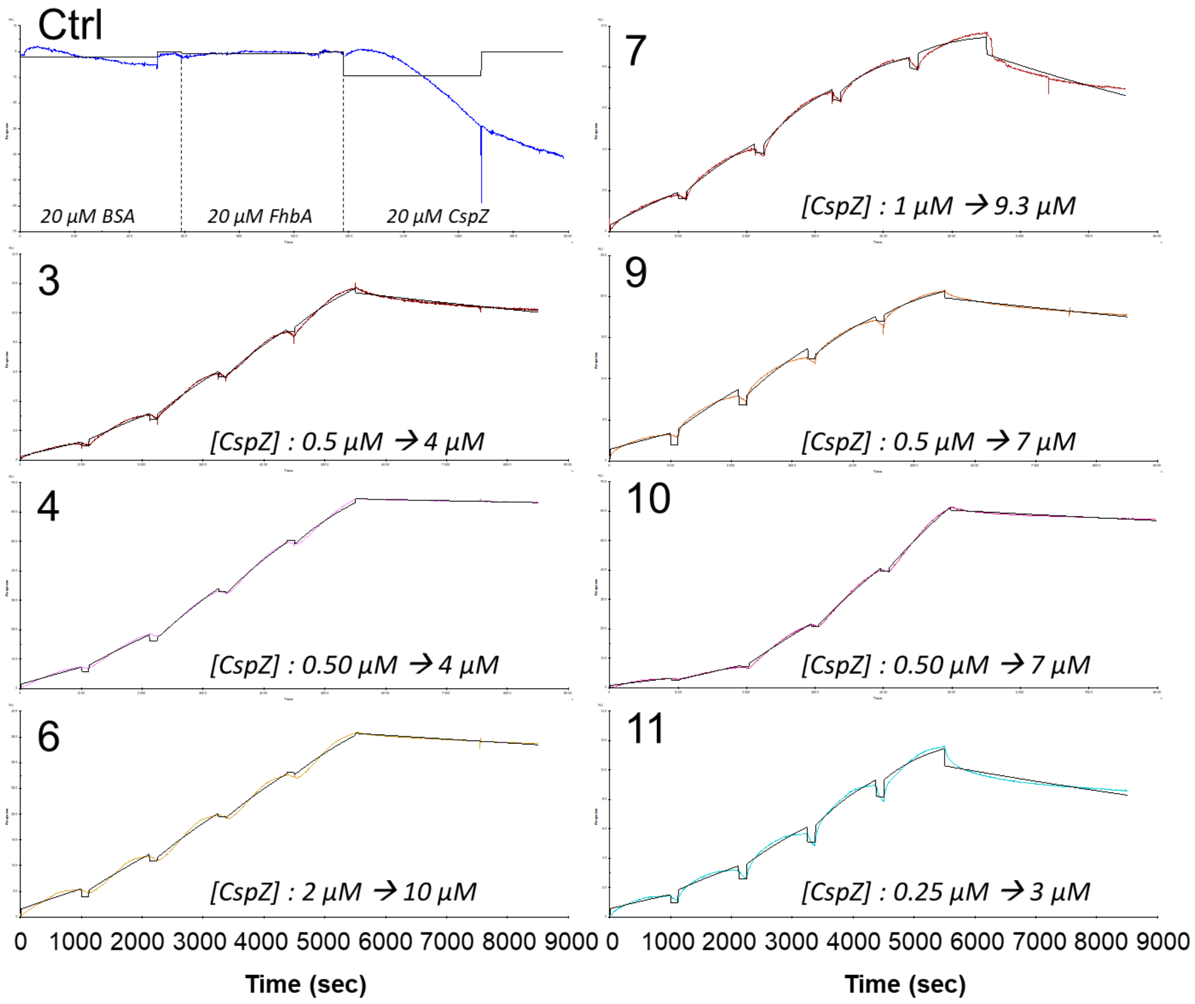


**Supplementary Figure 8 : SPR experiments for the selected aptamers as ligands, using a range of CspZ concentrations (from 0.25 µM to 20 µM depending of the aptamers) as analyte.**

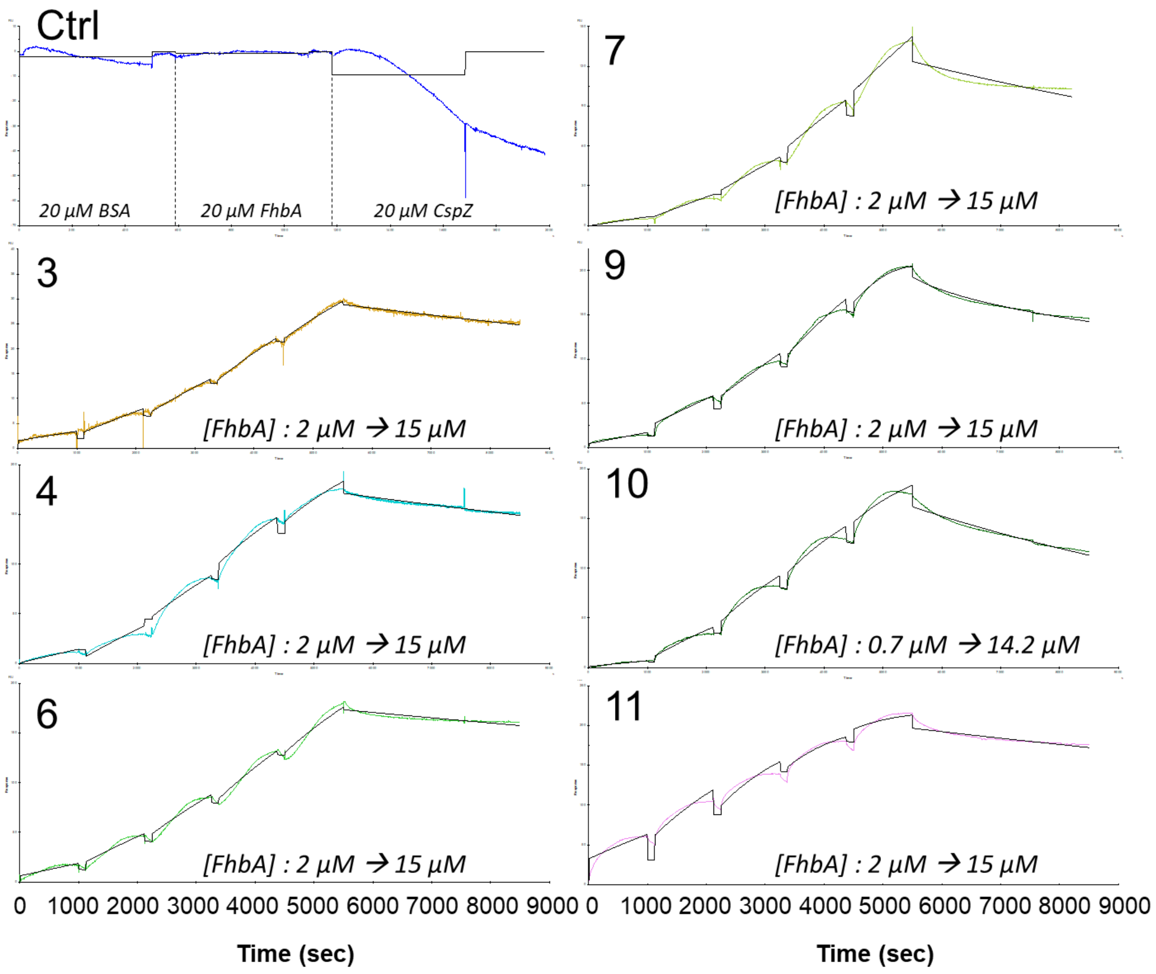


**Supplementary Figure 9 : SPR experiments for the selected aptamers as ligands, using a range of FhbA concentrations (from 0.7 µM to 20 µM depending of the aptamers) as analyte.**

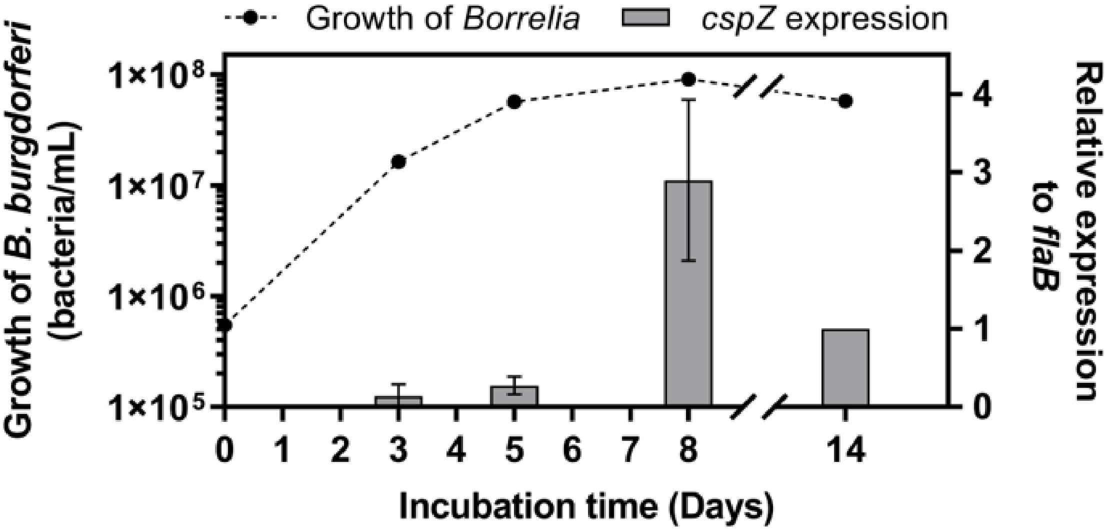


**Supplementary Figure 10 : Expression of *cspZ* during the growth of *B. burgdorferi* B31. The growth of *Bb* was performed in BSK-H at 33°C. Concentrations of spirochetes were determined by counting bacteria under a dark field microscope. *cspZ* expression levels were determined by RT-qPCR. Despite previous reports in the literature suggesting variation in *flaB* expression during cultivation[2], this study observed negligeable changes in *flaB* expression at t=14 days. Consequently, the *cspZ* expression levels were obtained by normalising to expression levels of the gene encoding *flaB* (internal control) at t=14 days. Data represent mean values ± standard deviation from three independent experiments.**

[2] W.K. Arnold, C.R. Savage, C.A. Brissette, J. Seshu, J. Livny, B. Stevenson, RNA-Seq of *Borrelia burgdorferi* in Multiple Phases of Growth Reveals Insights into the Dynamics of Gene Expression, Transcriptome Architecture, and Noncoding RNAs, PLOS ONE 11 (2016) e0164165. https://doi.org/10.1371/journal.pone.0164165.

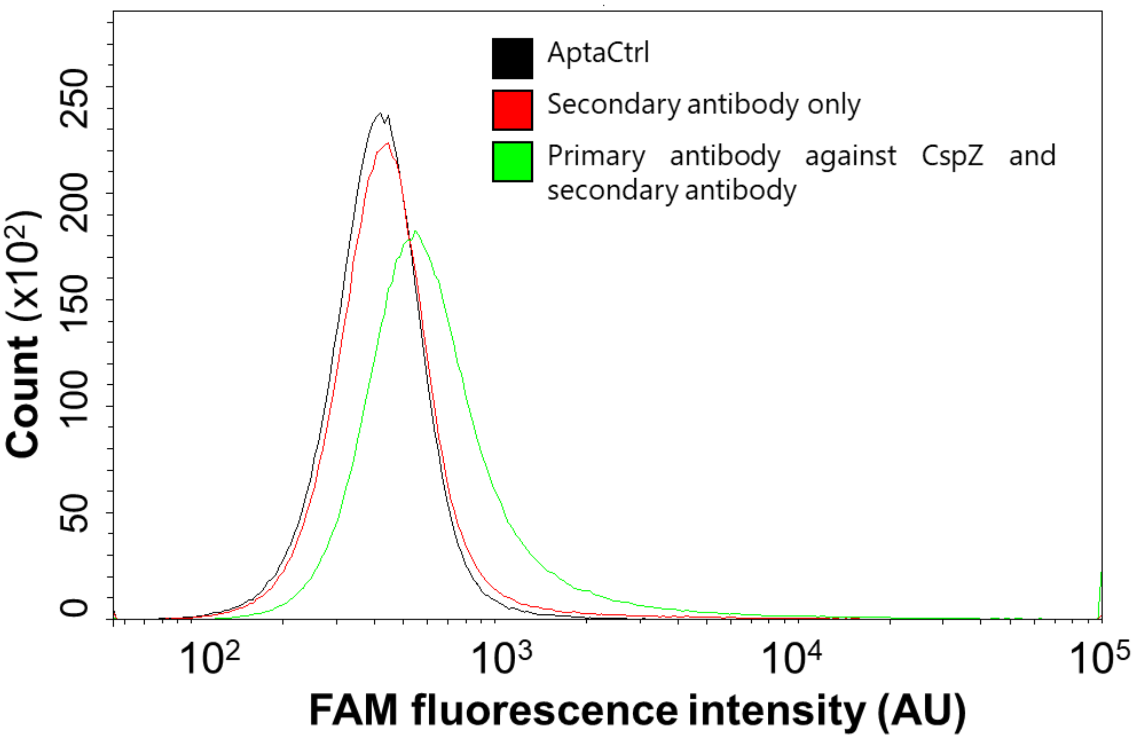


**Supplementary Figure 11 : Fluorescence of *Bb*ss induced by various controls using flow cytometry. The positive control (polyclonal antibody directed against CspZ revealed by a secondary antibody) is shown in green, the negative control (secondary antibody only) in red, and negative aptamer control AptaCtrl in black.**

**Supplementary Table 1: Sequence of the most representative aptamers, including a random sequence (AptaCtrl) used as the negative control for characterization studies. Each aptamer is flanked by constant 5' (ATACCAGCTTATTCAATT) and 3' (AGATAGTAAGTGCAATCT) regions.**

| **Name** | **Sequence of the random region (5'-3')** |
|---|---|
| Apta1 | CGAGGCCAGAAAAAGGGGAATGCGCCATAGCGTGTGGAAT |
| Apta2 | GTCAGAGGTGGAATGGCCGACTGAAGGGGAATGTCAGACA |
| Apta3 | GAACCGGGATGGGAGGGAGGGGGTGGAGGAGGCAGTTCAA |
| Apta4 | CACTTGGTGGTGGTGGCGGGATGGGATGGGTTGGGTTTGT |
| Apta5 | GTAACGTGAGGGAAACACTCGGTGCCTGTCATCCCTACAA |
| Apta6 | TGGGGCAAGGGAGGGCGGGGGCAGCGGCGGTACGAATTGA |
| Apta7 | AACGCGGATAGGGTGAGGGATGTTGGCGGTTTATGTGATA |
| Apta8 | CGGCATGTAGTCAACACAGCTGCTGGACGCGAGTATGCTT |
| Apta9 | TGGCAATGGAGGGGTGGTTGGAGGGGGGAATGTGGGGTT |
| Apta10 | TGGCAATGGGGGGTGGTCGGAGGGGGTTTACGTGGGTTCG |
| Apta11 | GGTTGGTCTGGTTGGCCCGTGTGTCATTACGGGTTGGATA |
| Apta12 | GGTGTGGTTTGGCACGTGGAACGTAGGGTGTGTCATTGAT |
| Apta13 | GCTCGGGTAGTAGCAAGGACCGTCTGAGGGTAATTAGCAG |
| AptaCtrl | CCCGGCCGACTTTGAACTTCAAGATAGGATCGACGTATGT |

**Supplementary Table 2 : Experimental conditions for each round of selection**

| Round | Input DNA (Folded ssDNA) | | CspZ-Beads complex | | | Ratio | Binding Time | Wash Number | Wash volume | ssDNA recovered after PCR | Monitor Index | | |
|---|---|---|---|---|---|---|---|---|---|---|---|---|---|
| | µg | pmol | Beads (µL) | Capture system | Target (nmol) | Ratio Protein/ ssDNA | (min) | - | (µL) | pmol | Output DNA (µg) | pmol | Output/ Input Ratio (%) |
| **Round 0 Depletion** | 122.6 | 5200 | 50<br>50 | Ni-NTA<br>GSH | - | - | 330 | - | - | | - | - | 92.3 |
| **1** | 113.5 | 4800 | 160 | Ni-NTA | 7.3 | 1.5 | 90 | 3 | 500 | 312 | 0.379 | 16.0 | 0.3 |
| **2** | 4.6 | 200 | 160 | Ni-NTA | 7.3 | 36.5 | 60 | 5 | 500 | 215 | 0.059 | 2.5 | 1.3 |
| **3** | 4.6 | 200 | 160 | Ni-NTA | 7.3 | 36.5 | 60 | 5 | 500 | 191 | 0.357 | 15.3 | 7.6 |
| **4** | 4.6 | 200 | 50 | GSH | 3.9 | 19.3 | 60 | 5 | 500 | 249 | 0.199 | 10.6 | 5.3 |
| **5** | 4.6 | 200 | 50 | GSH | 3.9 | 19.3 | 60 | 5 | 500 | 280 | 0.214 | 13.2 | 6.6 |
| **6** | 4.6 | 200 | 160 | Ni-NTA | 7.3 | 36.5 | 60 | 5 | 500 | 269 | 0.276 | 11.8 | 5.9 |
| **7** | 4.6 | 200 | 160 | Ni-NTA | 7.3 | 36.5 | 60 | 5 | 500 | 345 | 0.559 | 23.9 | 11.9 |
| **8** | 4.6 | 200 | 160 | Ni-NTA | 7.3 | 36.5 | 30 | 8 | 500 | 404 | 0.313 | 13.4 | 6.7 |
| **9** | 4.6 | 200 | 160 | Ni-NTA | 7.3 | 36.5 | 60 | 5 | 500 | 479 | 1.332 | 42.6 | 21.3 |
| **10 (Cell-SELEX)** | 4.6 | 200 | - | Bacteria *Bbss B31* | *10 mL culture* | - | 60 | 2 | 500 | 270 | 4.416 | 177.2 | 88.6 |
| **11** | 4.6 | 200 | 160 | Ni-NTA | 7.3 | 36.5 | 60 | 5 | 500 | 430 | 0.498 | 21.3 | 10.6 |
| **12** | 4.6 | 200 | 160 | Ni-NTA | 7.3 | 36.5 | 60 | 5 | 500 | 383 | 0.594 | 25.4 | 12.7 |

*Red* indicates the change in the experimental conditions

**Supplementary Table 3 : Nucleotide composition (%) of the random region after each round of selection**

| | A | C | G | T |
|---|---|---|---|---|
| **Library** | 19.07 | 20.37 | 32.53 | 28.02 |
| **Round 1** | 19.95 | 20.76 | 33.66 | 25.63 |
| **Round 2** | 18.74 | 22.38 | 31.66 | 27.23 |
| **Round 3** | 19.18 | 25.56 | 30.16 | 25.10 |
| **Round 4** | 19.25 | 25.20 | 29.87 | 25.67 |
| **Round 5** | 19.43 | 21.39 | 35.43 | 23.75 |
| **Round 6** | 23.16 | 24.11 | 30.51 | 22.22 |
| **Round 7** | 22.84 | 22.15 | 32.07 | 22.94 |
| **Round 8** | 22.84 | 18.11 | 35.74 | 23.31 |
| **Round 9** | 27.51 | 16.12 | 37.77 | 18.60 |
| **Round 10** | 20.33 | 13.53 | 46.09 | 20.05 |
| **Round 11** | 27.77 | 15.21 | 41.81 | 15.21 |
| **Round 12** | 28.97 | 15.86 | 40.16 | 15.00 |

**Supplementary Table 4 : Number of sequences obtained by NGS**

| | Initial number of reads | Trimmed sequences[1] | Sequences number after quality and length filter |
|---|---|---|---|
| **Library** | 125 556 | 120 355 | 105 772 |
| **Round 1** | 106 657 | 90 490 | 79 374 |
| **Round 2** | 96 756 | 87 209 | 74 824 |
| **Round 3** | 103 584 | 93 576 | 80 862 |
| **Round 4** | 105 396 | 89 673 | 76 090 |
| **Round 5** | 147 163 | 140 816 | 107 793 |
| **Round 6** | 179 556 | 161 509 | 141 575 |
| **Round 7** | 167 167 | 155 712 | 138 354 |
| **Round 8** | 159 569 | 148 181 | 132 343 |
| **Round 9** | 143 785 | 139 729 | 125 641 |
| **Round 10** | 138 709 | 134 306 | 113 638 |
| **Round 11** | 105 927 | 97 775 | 85 234 |
| **Round 12** | 139 492 | 135 692 | 119 736 |

[1] The initial trimming criteria was performed as described in Material and Methods section.
